# 1,000 ancient genomes uncover 10,000 years of natural selection in Europe

**DOI:** 10.1101/2022.08.24.505188

**Authors:** Megan K. Le, Olivia S. Smith, Ali Akbari, Arbel Harpak, David Reich, Vagheesh M. Narasimhan

**Author notes:** Co-corresponding authors **Corresponding authors** Correspondence to Vagheesh M. Narasimhan, David Reich and Arbel Harpak.

## Abstract

Ancient DNA has revolutionized our understanding of human population history. However, its potential to examine how rapid cultural evolution to new lifestyles may have driven biological adaptation has not been met, largely due to limited sample sizes. We assembled genome-wide data from 1,291 individuals from Europe over 10,000 years, providing a dataset that is large enough to resolve the timing of selection into the Neolithic, Bronze Age, and Historical periods. We identified 25 genetic loci with rapid changes in frequency during these periods, a majority of which were previously undetected. Signals specific to the Neolithic transition are associated with body weight, diet, and lipid metabolism-related phenotypes. They also include immune phenotypes, most notably a locus that confers immunity to *Salmonella* infection at a time when ancient *Salmonella* genomes have been shown to adapt to human hosts, thus providing a possible example of human-pathogen co-evolution. In the Bronze Age, selection signals are enriched near genes involved in pigmentation and immune-related traits, including at a key human protein interactor of SARS-CoV-2. Only in the Historical period do the selection candidates we detect largely mirror previously-reported signals, highlighting how the statistical power of previous studies was limited to the last few millennia. The Historical period also has multiple signals associated with vitamin D binding, providing evidence that lactase persistence may have been part of an oligogenic adaptation for efficient calcium uptake and challenging the theory that its adaptive value lies only in facilitating caloric supplementation during times of scarcity. Finally, we detect selection on complex traits in all three periods, including selection favoring variants that reduce body weight in the Neolithic. In the Historical period, we detect selection favoring variants that increase risk for cardiovascular disease plausibly reflecting selection for a more active inflammatory response that would have been adaptive in the face of increased infectious disease exposure. Our results provide an evolutionary rationale for the high prevalence of these deadly diseases in modern societies today and highlight the unique power of ancient DNA in elucidating biological change that accompanied the profound cultural transformations of recent human history.

## Main

Gene-culture co-evolution—whereby cultural adaptations including technological developments lead to new lifestyles that change selection pressures—have been widely discussed as a potential major driver of genetic adaptation^1^. To date, however, there have been few empirical examples, possibility due to the lack of ancient DNA data in sufficient sample sizes to reveal changes in allele frequencies before and after cultural change. This deficiency can be addressed with large ancient DNA datasets. Several central hypotheses have been put forward regarding how human cultural evolution may have driven human biological evolution^2^.

The first hypothesis relates to metabolic traits. The advent of agriculture induced a shift toward starch-rich and less diverse diets, which would be expected to lead to selection for loci that more effectively metabolize such diets and address their deficiencies of key nutrients^3^. Farming may have paradoxically also contributed to food scarcity. In times of plenty and food stability, population growth occurred at much faster rates than in the hunting and gathering period. However, these larger populations could also have been subject to periods of famine due to drought, agricultural disease outbreaks, or poor food distribution which might lead to additional selection for reduced caloric demand or more efficient energy metabolism.

The second hypothesis relates to gene-culture co-evolution associated with immunity. As humans began living in closer proximity to domesticated animals in the Neolithic, they would have been exposed to disease affecting those animals. In the Bronze Age and Historical periods, larger increases in population size as well as population movement occurred due to improved technology and mobility.

However, this would also have radically increased the opportunity for transmission of infectious disease and pressures on the immune system to more effectively combat them. The immune system has innate aspects associated with inflammatory processes and adaptive aspects associated with recognition of specific antigens. Making both these arms of the immune system more active can have deleterious consequences, for example a propensity to inflammatory processes such as atherosclerosis and autoimmune disease.

A third hypothesis relates to behavior. As population sizes became larger, societies became more complex, hierarchical, and inter-dependent. Selection could plausibly have occurred on genetic variation affecting traits such as individualism and sociability. This could plausibly have had impacts on neuro-psychiatric traits, including autism, schizophrenia, and bipolar disorder.

Ancient DNA provides time series data regarding human evolution, making it possible to directly study past selection by tracking allele frequency changes over time. Such data provides information about when and where selection occurred that cannot be obtained through analysis of present-day populations and should make it possible to study the hypotheses about gene culture co-evolution in practice. Until recently, the large sample sizes required to carry out these studies with high statistical precision have not been available. The earliest efforts to study natural selection using ancient DNA data have therefore been limited^4–6^, often focusing on candidate loci or single traits^7–10^. More recent approaches have looked at selection genome-wide but focus on obtaining evidence of selection across the full range of time from the Paleolithic leading to modern Europeans^11–13^. Such analyses may miss out on selective events that might be operating only for short bursts in pre-history in response to cultural change. Some other approaches look at specific time slices in the data but require comparisons with simulations of demographic models that might not always be available for ancient genomes^14^. Other approaches utilize haplotype approaches that are unable to precisely identify the targets of selection^14,15^.

Here, to examine selection acting across several time intervals in human history, we assembled a large sample-size time transect from Holocene Europe comprising published data generated using the same technology that has been the source of more than 70% of published ancient DNA data to date: in-solution enrichment for about 1.2 million single nucleotide polymorphisms (SNPs). Studying this period and geographical region is interesting not only from the limited perspective of this place and time, but also for understanding the processes of natural selection over ten millennia of profound change in human lifestyle. These include the transition from hunting and gathering to farming, which resulted in major changes in diet as well as increased population density and proximity to animals. This period also includes the transition to state-level societies facilitated by metal-working, which led to large population densities, long-distance exchange of goods, and division of labor. Several ancient DNA studies have also sequenced bacterial and viral pathogens that caused epidemics in the last few millennia, including smallpox, the black death, and tuberculosis, suggesting that studying ancient DNA in a time transect might provide insights into human adaptation to these new infectious diseases^16–18^.

The complicated demographic history of human populations, which includes migration and mixture with neighbors, makes it challenging to determine whether natural selection or population mixture is the driving force behind changes in allele frequencies that occurred in the past. However, in Europe, multiple ancient DNA studies have provided excellent models for demographic history^4^. Here, we identify individual genetic loci as well as sets of alleles whose changes in frequency are inconsistent with the expectation under neutral evolution and these demographic models, and are therefore suggestive of selection. Given the large sample sizes spanning this time transect that provide a nearly gapless record of human populations in Europe in the Holocene, we are further able to estimate the timing of selection and generate hypotheses about its correspondence with major demographic and cultural changes.

### A time transect through Holocene Europe

We assembled genome-wide data from a total of 1,291 individuals from Holocene Europe dated to between 13,000 and 1,000 years before present (BP) (<u>Supplementary Table 1</u>). We restricted to individuals with at least 15,000 SNPs^19,20^. We only included unrelated (up to the third degree) individuals without significant contamination as assessed on the mtDNA or, in males, the X chromosome. We chose to only analyze data from libraries that were treated with the enzyme uracil-DNA glycosylase (UDG) prior to library preparation, which reduces characteristic cytosine-to-thymine errors associated with ancient DNA data, and that were then enriched in-solution at about 1.2 million SNP positions. For population history analysis, we generated pseudo-haploid calls at every location. For natural selection analysis, we retained read counts of the reference and alternate allele at every site for our likelihood calculation of allele frequencies (<u>Methods</u>). To be conservative and avoid false-positive signals of selection, we did not impute genotypes at untargeted positions due to potential biases associated with using a modern reference panel to phase and impute ancient genomes that are of low coverage (median coverage ∼0.9x) and could have different haplotype structure^21^. To avoid additional biases associated with misestimating allele frequencies with heterogeneous data, we did not include ancient shotgun data or modern data in our analysis.

We leveraged a model of the demographic history of Europe over the past 10,000 years that has been inferred from ancient DNA studies^4,5,22–29^. Broadly, these studies conclude that most Europeans in this period derive the great majority of their ancestry from three primary ancestral sources which came together in the course of two major demographic transitions corresponding to significant shifts in the archeological record. The first is the transition from hunting and gathering to farming, which was accompanied by a major population transition in central and western Europe. In this period, ancestry from the Mesolithic inhabitants of central and western Europe was largely displaced by ancestry from farmers whose ancestors originated in Anatolia, and who were amongst the first peoples in the world to use agriculture a few thousand years before. This economic and demographic shift began in southeastern Europe after around 6,500 BCE but had spread to the far reaches of Europe as well as Britain by 4,000 BCE. The second major demographic transition occurred during the shift from the Neolithic to the Bronze Age with the arrival of Steppe Pastoralists from the Eurasian Steppe. In the subsequent millennia leading to the Historical period, there were subtle shifts in the proportion of Steppe ancestry that largely arose from the homogenizing of populations with different Steppe ancestry proportions.

We assigned individuals to different groupings based on *f*_*4*_-statistics, time period (based on direct radiocarbon dates or well understood archaeological contexts), and geographic location. We removed individuals that were outliers from each time period and were found to have atypical ancestry of that period based on *f*_*4*_-statistics. The groupings of individuals were:

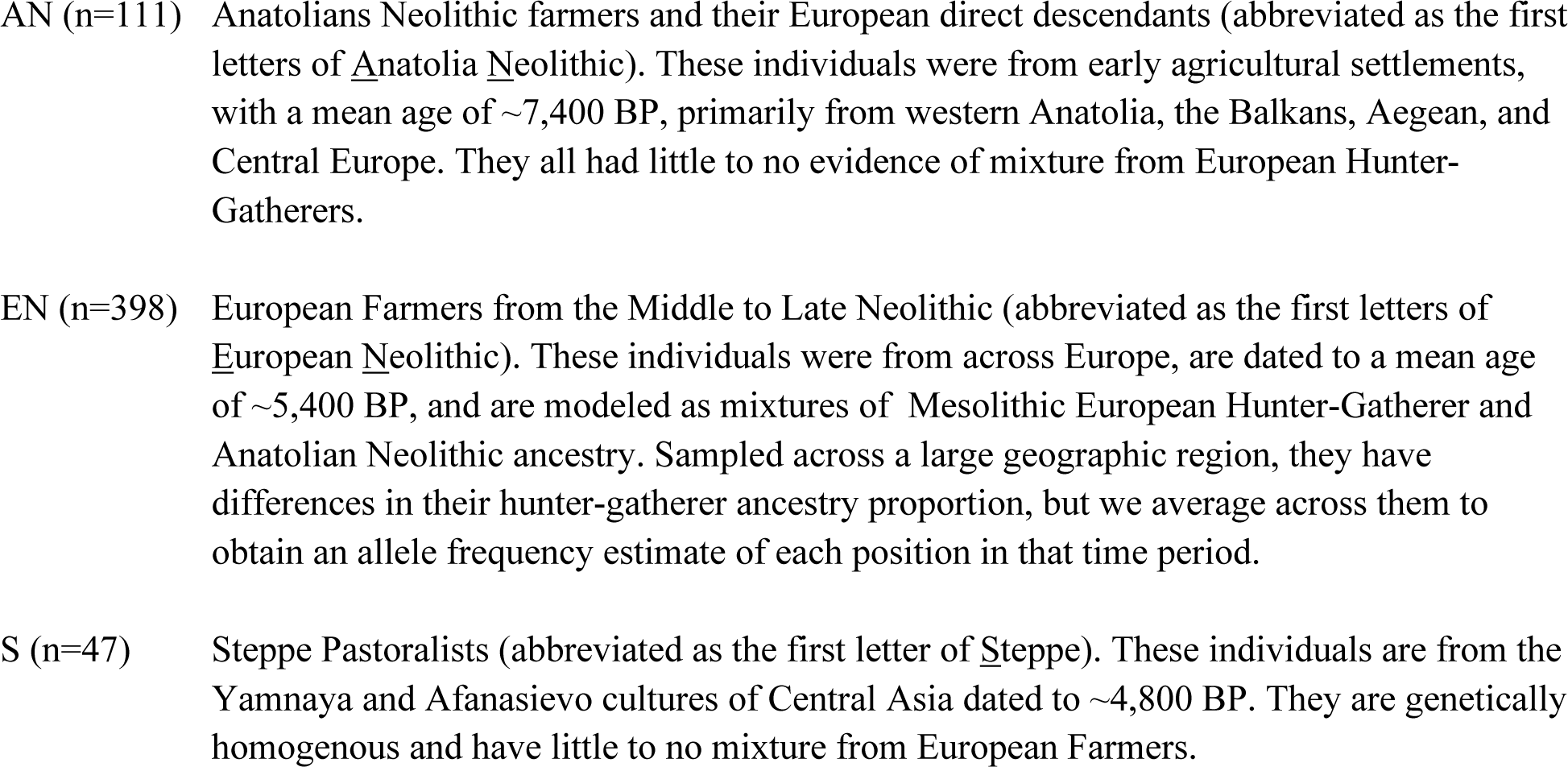
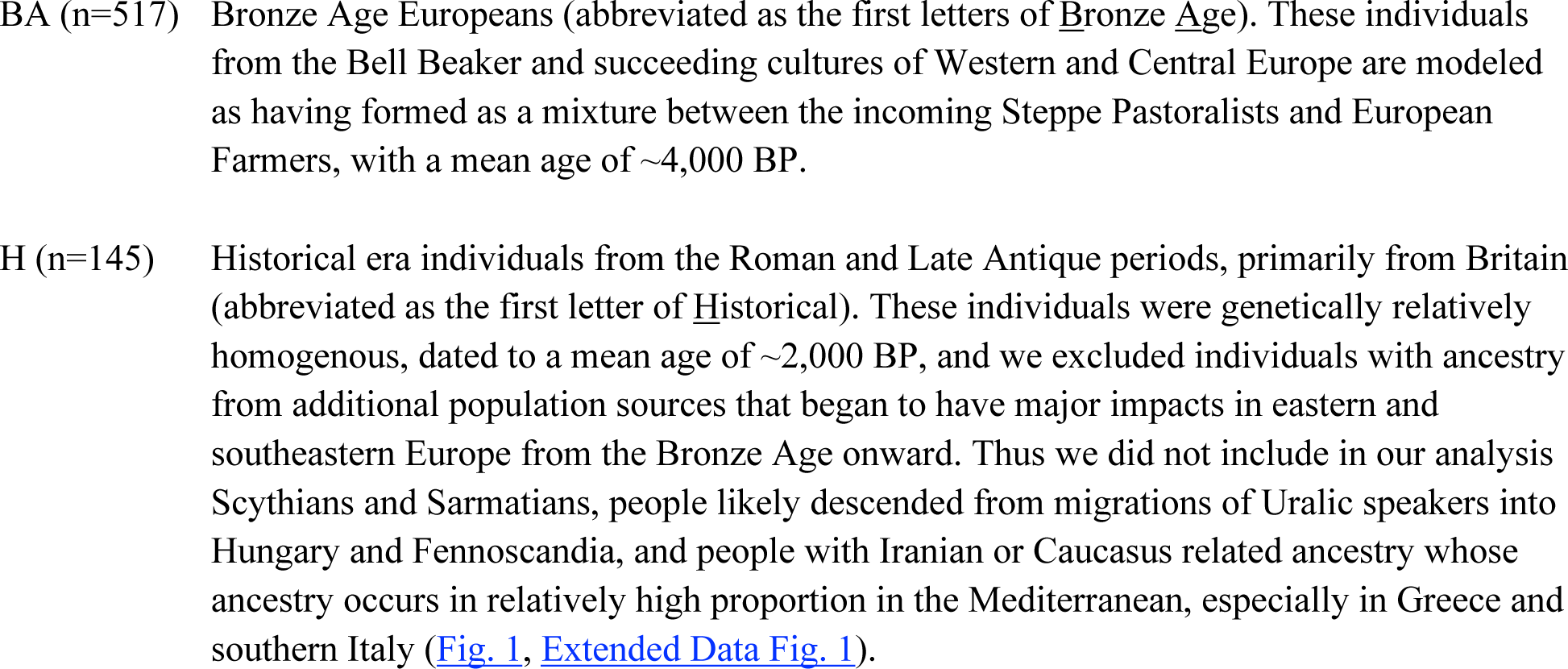

**Fig. 1:**
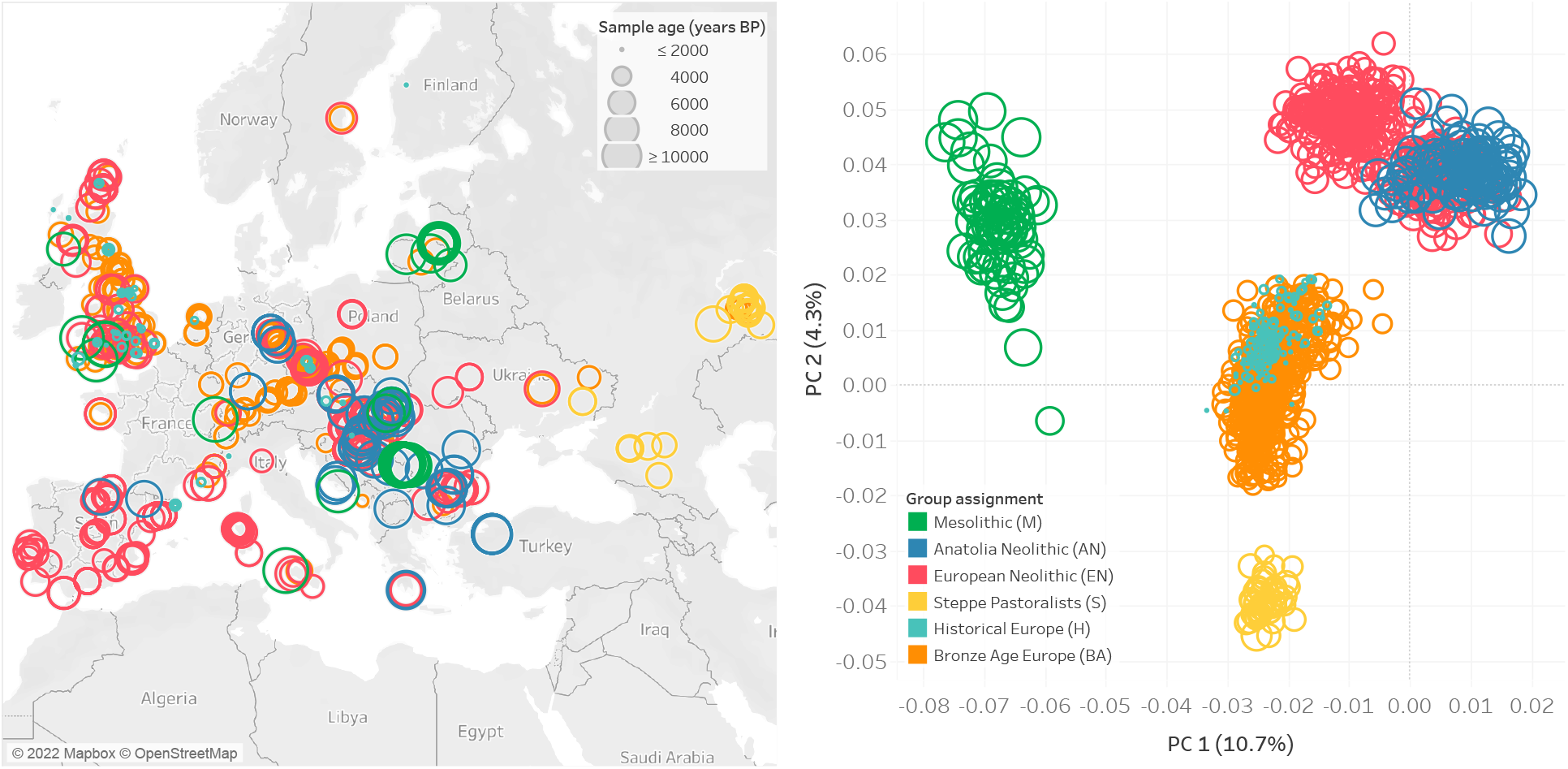
Geographic and temporal distribution of analyzed individuals. **a**, Geographic locations and group assignments (in color) for all individuals along with sample age represented by the size of the circular points. **b**, Principal Components Analysis of samples with the same grouping and coloring scheme as in **a**.

Full lists of all individuals, their assignments, and additional metadata can be found in <u>Supplementary Table 1</u>.

To model these demographic changes with our combined dataset, we used *qpAdm*, which evaluates demographic fit of a target population to various source populations genome wide and then estimates proportions of ancestry for each source^22^. We divided the roughly 10,000 year period into three non-overlapping time epochs, each of which spans just over 3,000 years: (1) the transition from hunting and gathering to farming (the Neolithic period), (2) the transition to the Bronze Age, and (3) the transition to large-scale state-level societies during the Historical period. To capture the major sources of admixture in epoch (1), we modeled European Farmers (EN) as a 16:84% mixture of European Hunter-Gatherers and Anatolian Farmers. For epoch (2), we modeled Bronze Age Europeans as a 48:52% mixture of European Farmers and Steppe Pastoralists. For epoch (3), we modeled Historical European samples as a 15:85% mixture of Bronze Age Europeans and earlier Neolithic Farmers (Fig. 2).

**Fig. 2:**
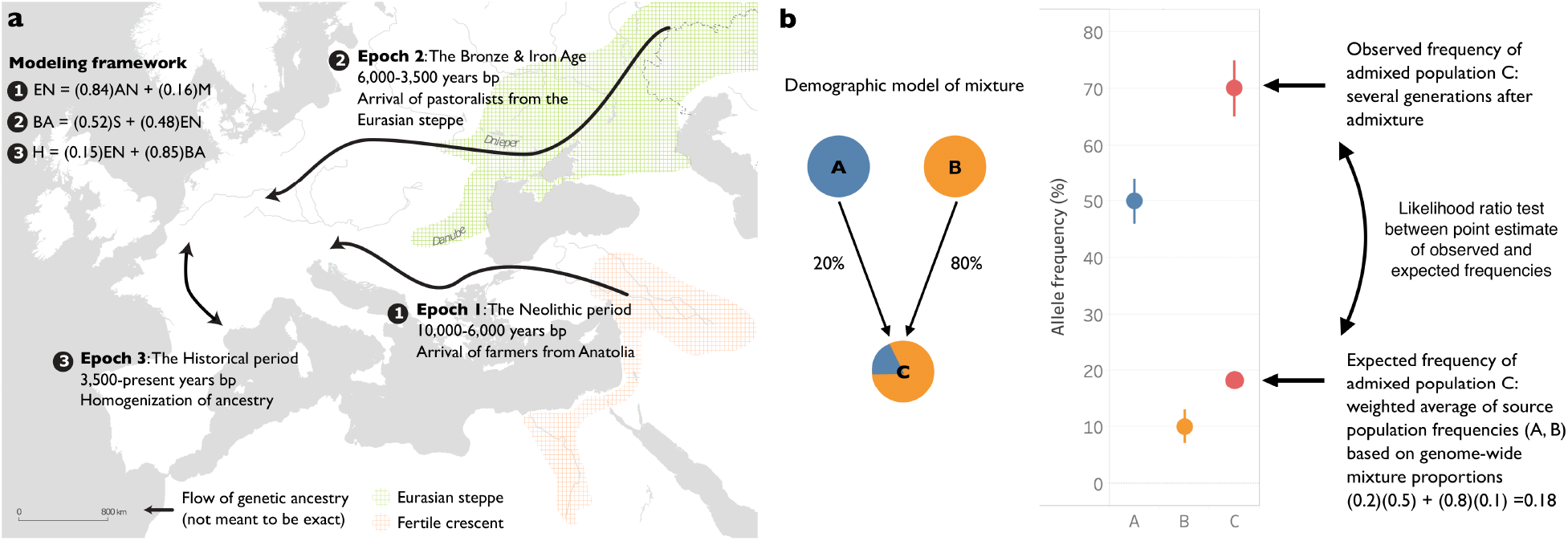
Description of approach to detect selection. **a**, Demographic changes in Europe over the past 10,000 years are driven by admixture between various populations across three major time epochs/periods. **b**, Visual description of our methodology. Under neutrality, the expected frequency of an allele is the weighted average of the source population frequencies. Large deviations from this genome-wide expectation can be identified as evidence for selection. In this case, the frequency of the allele in population C has risen to a frequency that is ∼50% above the expected frequency based on mixture proportions and suggests that natural selection has elevated the frequency of this allele in the time period since admixture.

### A scan for selection at individual loci

To identify candidate selected loci in our dataset, we used our three epoch model and applied a method that utilizes the admixture events that occurred in each epoch. Under neutrality, the allele frequency of an admixed population is expected to be the weighted average of the allele frequencies in the source populations that contributed to the admixture. Significant deviations from this expectation suggest that natural selection has acted at a particular locus (Fig. 2). After correcting for inflation of the test statistic independently in each of the three epochs, we used a cutoff of 5×10^−8^ as a genome-wide significance threshold. This is a common significance threshold in genome-wide association studies (GWAS)^30^, and also roughly corresponds to a *P* value of <0.05 after Bonferroni correction for a 1.2 million SNP target set (<u>Methods</u>). Previous work has examined the impact of sample size, the strength of selection, the time that selection has acted, mis-specification of the mixture proportions, and additional unmodeled mixtures in detecting selection using this method and has shown that, after applying a correction for genomic inflation, these issues result in reduced power but not an increased rate of false-positives^4,31^. Additional work on the same method with slightly different statistical formulation has confirmed this robustness to deviations from the model^32^. To further study the effect of model misspecification as well as the effect of sample size on our power to detect signals, we carried out two additional analyses. First, we examined our model’s robustness to mis-estimates of the admixture proportions and found that deviations on the order of 15% resulted in little reduction in power (Extended Data Fig. 2). Second, we found that reduced sample size below 80% of the dataset size used for analysis has a major effect on power to detect selection signals (Extended Data Fig. 3).

Following a previous strategy used to mitigate false-positives in ancient DNA scans of selection due to biases affecting the sequences aligning to a particular variant, we considered loci to be candidates for selection if at least 3 alleles within 1 Mb of each other and the causal gene significantly deviated from their expected frequency^4^. This distance is also in agreement with a recent study examining the optimal window size for linking GWAS-associated SNPs to causal genes^33^. To determine if functional categories of genes were significantly associated with selection signals, we carried out enrichment analysis using FUMA^34^, which maps SNPs to genes and performs gene set enrichment analysis for GWAS and GO annotations incorporating LD information as well as gene matching by length and conservation scores (Methods).

### 25 time-resolved candidate signals of natural selection

Across all epochs, we discovered a total of 25 regions containing alleles with frequency changes that significantly deviated from genome-wide expectation (Fig. 3, Extended Data Fig. 4, Extended Data Fig. 5, Extended Data Fig. 6, Table 1). The only locus that contained alleles with significant evidence of selection across all time periods was the Major Histocompatibility Complex (HLA) on chromosome 6, which encodes cell surface proteins that are a critical part of the human adaptive immune response.

**Table 1:**
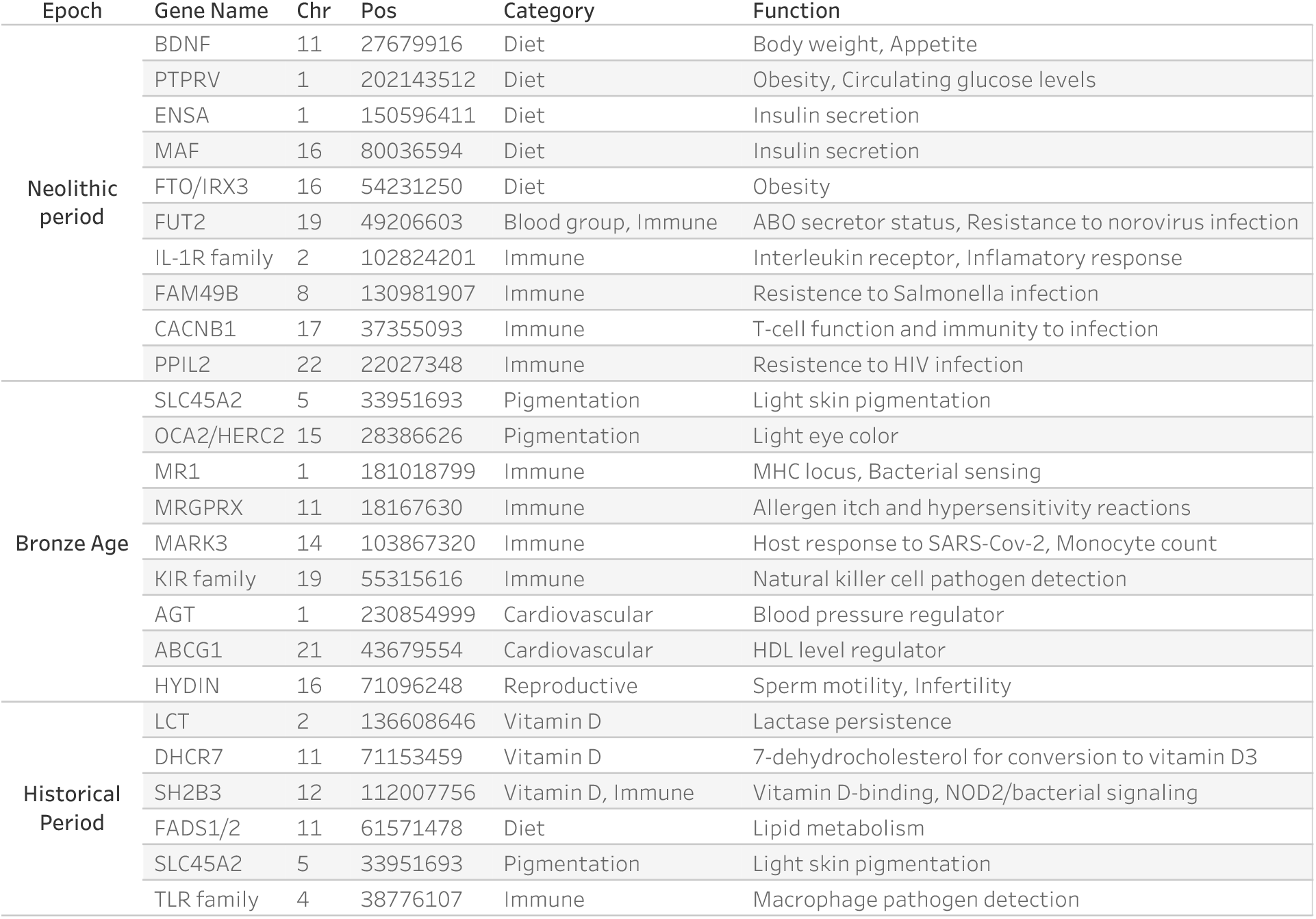
Summary of genes with evidence of selection during the three epochs in Europe. The HLA region, which appears to be under selection in all epochs, is not shown.

**Fig. 3:**
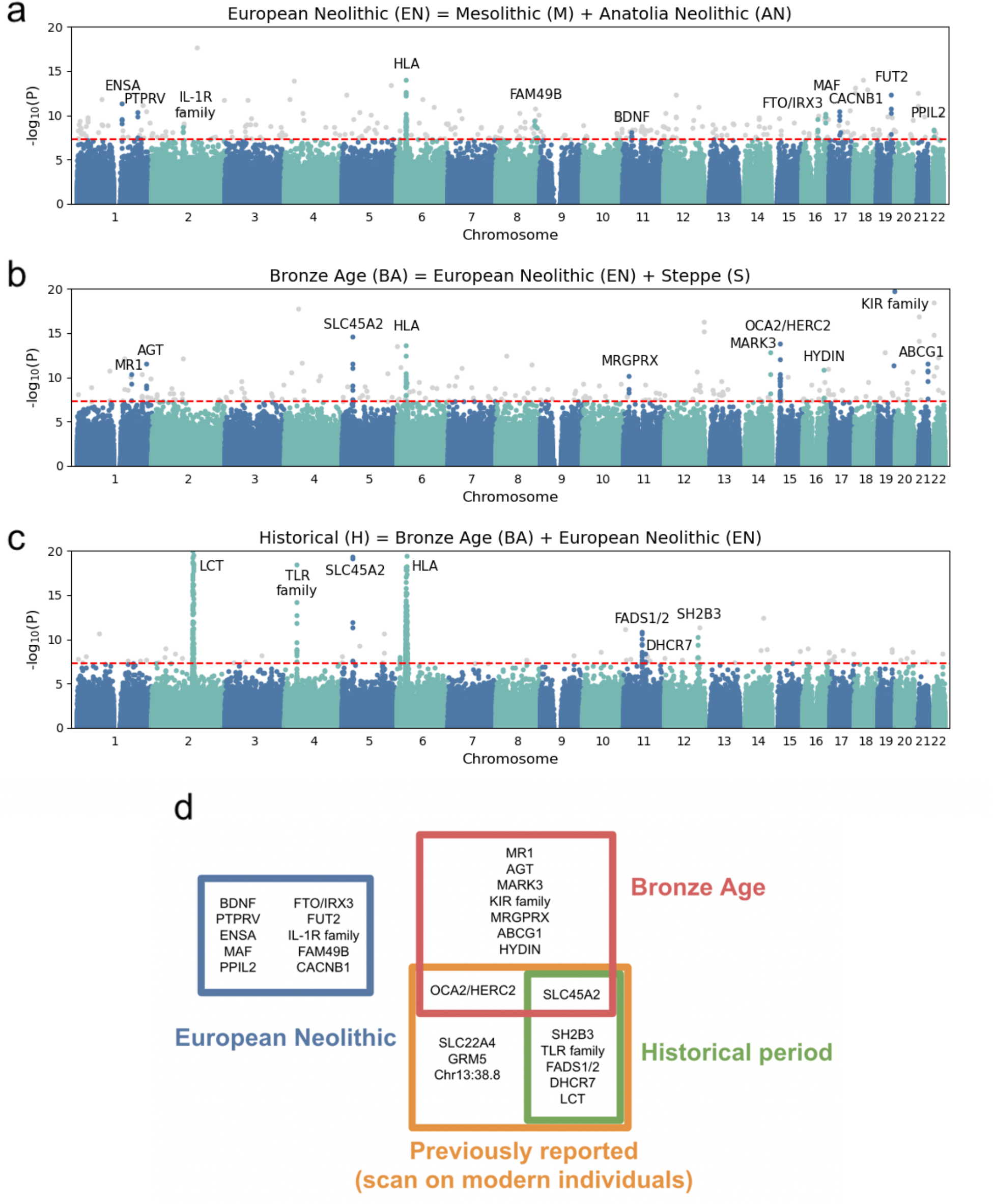
Signals of natural selection in three epochs. **a-c**, Manhattan plots of *P* values for the likelihood ratio test for selection (<u>Methods</u>; Fig. 2) in the Neolithic, Bronze Age and Historical period. The red line shows the genome-wide significance threshold (5 × 10^−8^). **d**, Venn diagram showing the overlap of variants seen in each epoch and the variants that were previously published (Mathieson et al.^4^) from a scan on present day humans. HLA, which was previously published and seen in all epochs, is not shown.

### Candidate selective signals that were most intense during the early phases of the transition to agriculture

In the first epoch representing the transition from hunting and gathering to farming, we discovered individual signals that were plausibly associated with a transition to a high starch, carbohydrate heavy diet, to which the genomes of the two ancestral populations were not yet fully adapted.

First, we observed several alleles at the *FTO/IRX3* locus, the locus that has the largest effect on predisposition to obesity in humans^35^. Reduction in gene expression of this gene has shown 30% reduction in weight in humans and model organisms^36^. The region and variants that were significant in our scan are in the promoter region of the *IRX3* gene and are in high LD with variants that are expression quantitative trait loci (eQTLs) in human adipose/subcutaneous tissues reported by the GTeX consortium^37^. *IRX3* expression is known to increase body weight^36^, and variants that decrease the expression of *IRX3* increased in the Neolithic transition, which suggests that there may have been selection for reduced body weight specifically during this time.

We also found alleles that were significant in the gene *PTPRV*. Mice homozygous for a knock-out allele at this gene exhibit increased resistance to diet-induced obesity and decreased circulating glucose levels^38^. Other studies have shown that *PTPRV* also contributes to *FTO*’s role in adipogenesis with simultaneous knockdown of both genes restoring adipogenesis activity that is lost when just *FTO* alone was knocked down^39^. Selection for these variants affecting adipogenesis could be adaptive in the course of an economic transition between a hunting and gathering lifestyle to a farming-based lifestyle, which would have involved a greater reliance on starch-based diets and different patterns of feast-and-famine.

We detected another candidate in the gene *ENSA*, which acts as a stimulator of insulin secretion by interacting with the protein encoded by *ABCC8*, a sulfonylurea receptor which plays a key role in the control of insulin release in pancreatic beta cells. We also observed variants with likely similar function in the regulatory region of the gene *MAF* (rs4073089) which promotes pancreatic development and regulates insulin gene transcription^40^. Another candidate of selection is on the missense variant rs6265 that occurs at around 19% frequency in modern Europeans on the *BDNF* (Brain-derived neutrophil factor) gene, which has been associated with regulation of body weight and has a mechanistic role in Type 2-diabetes in humans and model organisms^41,42^.

Several signals of selection in this period are associated with immune-related functions. We detect a signal in the gene *FUT2*, the human secretor locus that encodes α(1,2)-fucosyltransferase, and determines the secretion status of the ABO blood group antigens. Individuals homozygous for the *FUT2* non-secretor genotype are resistant to infection with norovirus^43^, suggesting that individuals homozygous for non-secretor status may be unable to mediate host-microbe interactions. The variants that are significant in *FUT2* have also been associated with plasma B12 levels^44^, a vitamin that is largely unavailable from plant-based food sources—in particular, it is virtually absent in wheat and barley, which make up the bulk of the Neolithic agricultural package—but plentiful in animal products.

Another significant signal is at the Interleukin 1 receptor, type II (*IL-1R2)* which is expressed on lymphoid and myeloid cells including monocytes, neutrophils, macrophages, B, and T cells, and has been implicated as a susceptibility locus for a number of autoimmune diseases^45^. We also found signals of selection at alleles in the gene *PPIL2* which encodes a cyclophillin, a class of proteins that bind to ciclosporin (cyclosporin A), an immunosuppressant which is used to suppress rejection after internal organ transplants. *PPIL2* and other cyclophillins are also recruited by the Gag polyprotein during HIV-1 infection, and its incorporation into new virus particles is essential for HIV-1 infectivity^46^, suggesting that selection at this locus may reflect selection against HIV-like retroviruses. Another gene that is under selection is *CACNB1*, a regulator of T-cell function. Mice lacking in the *CACNB1* gene have been shown to be immune-deficient to viral infection^47^.

Finally, we detect evidence for selection at variants in *FAM49B. In-vitro* as well as *in-vivo* studies have recently shown this gene is a T-cell regulator and confers host resistance to *Salmonella* infection^48,49^. *FAM49B* negatively regulates *RAC1* signaling, thereby attenuating processes such as macropinocytosis, phagocytosis, and cell migration. This enables the protein to counteract *Salmonella* at various stages of infection, including bacterial entry into non-phagocytic and phagocytic cells as well as phagocyte-mediated bacterial dissemination^48^. Evidence for *Salmonella enterica* adaptation to the human host through rise of specific pathogenic mutations has been detected through bacterial sequencing of ancient DNA time transects from Europe and has been timed to the Neolithic period^50^. Our observation of candidate regions of selection in humans (the host) at this locus during the same time period is compatible with human-pathogen coevolution at a time of major cultural change.

Consistent with the associated gene function of individual variants, we found an enrichment of candidates near genes related to fatty acid metabolism and digestion, serum metabolite levels, and diseases of the digestive system such as Crohn’s disease and ulcerative colitis (<u>Supplementary Table 2</u>).

### Candidate selective signals most intense during the transition to the Bronze Age

In the Bronze Age, we do not detect evidence for continued selection on candidate variants that are directly associated with a change in diet. Instead, we found evidence for selection at or near genes that affect skin and eye pigmentation.

The strongest signal is at the allele rs16891982, in the gene *SLC45A2*, which is known to play a major role in light skin pigmentation, and for which there has been previous evidence for selection^51^. The second strongest signal based on our analysis is in the allele rs11636232 in *OCA2/HERC2*, which is a primary determinant of light eye color in Europeans^4,52^.

As in the Neolithic period, we also found several candidate genes involved in immunity beyond those seen in the HLA region. We observed selection at rs10797666 in the major histocompatibility complex class I-related gene *MR1*, which is an immune sensor of microbial ligands, including *Mycobacterium tuberculosis, Streptococcus pyogenes, Salmonella enterica* and *Escherichica coli*^53^. We also find evidence of selection at a number of alleles in genes in the killer-cell immunoglobulin-like receptor locus (*KIR* gene family), which are expressed on the cell membrane of natural killer cells. *KIR* receptors interact with major histocompatibility molecules to detect pathogen-infected cells and have a crucial role in host defense. This locus is highly polymorphic across human populations worldwide, and diversity at this locus has been correlated with pathogen load across populations^54^. We also find evidence for selection at the *MRGPRX3-4* locus, which includes genes that are key physiological and pathological mediators of itch and related mast cell-mediated hypersensitivity reactions, as well as potential targets to reduce drug-induced adverse reactions^55,56^. A final immune-related candidate is the gene *MARK3*, which is a host protein that is one of the key interactors with SARS-CoV-2 and is important in mediating the maladaptive host response to COVID-19. The allele under selection appears to be linked to a lead signal for monocyte count, which has now been shown to be important in the pathology of COVID-19^57–59^.

There is direct ancient DNA evidence for pathogen infection being a major source of mortality in the Bronze Age. The earliest evidence for *Yersinia pestis* infections in Europe ascertained from ancient DNA comes from the Bronze Age at times particularly close to or after the arrival of pastoralists from the Eurasian Steppe, from where both of these pathogens have been recovered from humans several millennia prior to their first evidence in Europe^60,61^. Thus, pathogens entering Europe along with Steppe Pastoralists in the Bronze Age could have been a driving force behind changes in these immune related genes.

We also observed significant frequency deviation from the expectation due to genetic drift in alleles lying on several genes related to cardiovascular disease. One candidate is in the angiotensin gene, *AGT*, which causes vasoconstriction and increases blood pressure^62^. The protein encoded by AGT is a frequent antagonist in drugs that treat heart disease. Additionally, we also observed another locus that reached significance, rs915843, which is a missense allele in *ABCG1*, a gene that controls tissue lipid levels and the efflux of cellular cholesterol to HDL^63^.

Finally, we observed candidates in genes where mutations have been linked to reproduction. One of our significant variants is at rs7188473, a splice mutation in the gene *HYDIN*. Homozygous carriers of this allele suffer from primary ciliary dyskinesia-5, which affects sperm motility and leads to male infertility^64^.

More broadly, across this epoch, we find a statistically significant enrichment of signals near genes related to skin, hair, and eye pigmentation (<u>Supplementary Table 3</u>).

### Candidate selective signals most intense during the transition to the Historical period

The variant with the strongest significant deviation from expectation is in the *LCT* gene, which is responsible for conferring the ability to digest lactose in adulthood in Europeans. This is also consistently the strongest signal of natural selection detected in scans in modern Europeans, and in line with findings in previous publications^65^, this allele appears to have experienced its major change in frequency primarily in the past few thousand years, and not during the Bronze Age when the allele was first introduced in central and western Europe.

We found a selective signal in *DHCR7* (the focal SNP that deviates most from expectation is in an enhancer region several kb upstream of the gene), which governs availability of 7-dehydrocholesterol for conversion to vitamin D3 by the action of sunlight on the skin. Milk is rich in 7-dehydrocholesterol, suggesting that selection on this locus as well as *LCT* might have been related to the need for increased production of vitamin D^66^. This locus has also been linked to several auto-immune diseases. We also detect evidence for deviation in allele frequency from expectation in the missense variant rs653178 in the gene *SH2B3*. This allele doubled in frequency from the Bronze Age to the Historical period and is a major risk locus for Celiac disease. The allele we identify as a signal of selection has recently also been shown to be fine-mapped in a GWAS for vitamin D binding^67^. Functional investigation of the effect of the *SH2B3* genotype in response to lipopolysaccharide and muramyl dipeptide revealed that carriers of the *SH2B3* allele showed stronger activation of the *NOD2* recognition pathway. This suggests that *SH2B3* also plays a role in protection against bacterial infection^68^.

The second strongest signal was in the gene encoding the toll-like receptor locus *TLR*, which is expressed on the membranes of leukocytes. Variants at this locus have been associated with host immune response against a variety of diseases, including *Heliobacter pylori* infection, leprosy, plague, and tuberculosis.

We detect continued change in frequencies of variants at the *SLC45A2* gene, driving the light-pigmentation allele under selection to near fixation in Historical samples.

Finally, we found candidate variants on the genes *FADS1* and *FADS2*, which are involved in fatty acid metabolism. Variation at this locus is associated with plasma lipid and fatty acid concentration. The most significant SNP (rs174550) at this locus has also been associated with decreased triglyceride levels, and perhaps selection at this locus could reflect transition to a starch-heavy diet. This locus has also experienced independent selection in non-European populations and is likely to be a critical component of adaptation to different diets. In agreement with previous work suggesting that natural selection acted on these alleles only after the Neolithic transition^7^, in our analysis we see that the major signal of selection at this locus is focused on the most recent epoch.

In this period we also find statistically significant enrichment in gene sets involved in a large variety of traits, from anthropometric traits such as BMI and autoimmune disease like Crohn’s and ulcerative colitis, to hormone-related disorders like hyperthyroidism, blood biomarkers such as serum metabolite and cholesterol levels, and cardiovascular disease traits (<u>Supplementary Table 4</u>).

### Timing of selection of alleles

By separating our analysis into different epochs, we were able to examine overlaps in candidate alleles across epochs as well as a previous scan examining modern Europeans from the 1000 Genomes Project. Outside of the HLA region, we found no overlaps of any of the loci discovered in the Neolithic period with any of the other epochs. All of the candidates we discovered in the Historical period had also been discovered in a scan comparing modern Europeans to ancient Europeans^4^. As expected, this scan was largely blind to selection during the European Neolithic, showing the value of direct comparison of groups of ancient DNA samples to study the selection that occurred in this time (Fig. 3).

### Polygenic selection

Evidence from contemporary genomes suggests that in recent human history, monogenic selective sweeps are rare^69,70^. Further, theoretical arguments and some empirical evidence in the last decade suggest that polygenic adaptation may be the more frequent mode of selection52,71–74. Therefore, to complement the picture we obtain of selection in the last 10,000 years from the monogenic genome scan, we sought to test for polygenic selection on complex traits. We did so by integrating the same signal of deviation from expected admixture proportions with trait-associations from genome-wide association studies (GWAS).

### Mitigating confounders of our tests for polygenic selection

Despite the theoretical appeal of screens for polygenic selections, clear evidence for polygenic selection has been elusive due to challenges in application and interpretation^71,72,75–79^. Here, we take an approach that offers more robustness against a major challenge: the portability of GWAS associations from contemporary GWAS to ancient samples.

GWAS-based estimates may often be poor or even biased with respect to individuals in populations different from the groups in which the GWAS was carried out, due to differences in ancestry, environment, or other characteristics. This can lead to biased or uninterpretable results^80,81^. This also applies for porting modern GWAS associations to ancient genomes. There are several reasons for poor portability. A major problem, which can lead to systematic biases, is uncorrected population stratification (axes of ancestry that correlate with a trait) in GWAS. Regardless of the cause of the correlation with the trait, numerous alleles that tag these ancestry axes may still associate with the trait. This problem may be amplified as GWAS sample sizes increase and many small effects become more statistically significant^75,76,82^.

We took two measures that are expected to reduce statistical power but increase robustness to population stratification. First, for our primary analysis we use GWAS summary statistics (for 38 case-control and 177 quantitative traits) from the Biobank of Japan (BBJ), rather than summary statistics based on GWAS in Europeans with higher sample sizes. Since West Eurasians and East Eurasians largely split from a common ancestral population more than about 40,000 years ago, the population structure present in the BBJ sample is expected to be uncorrelated with the main axes of population structure among Europeans. In addition, as suggested by Chen et al.^83^, linkage disequilibrium (LD) and allele frequency differences between the BBJ sample and the different ancient European target samples are mediated through a common ancestral population and thus should be approximately equal.

Following Chen et al.^83^, we evaluated residual stratification by examining the association between effect sizes estimated from each GWAS panel and PC loadings conducted on a set of diverse West Eurasian populations that reflects the various ancestry sources in Europe^23^. The first PC separates Steppe Pastoralists from Western European Hunter Gatherers, and the second PC separates Anatolian and Iranian Farmers from Steppe Pastoralists and Western European Hunter-Gatherers. To measure the impact of uncorrected stratification on estimated effect sizes for a set of ascertained trait-associated variants, we regressed the SNP effect sizes on the PC loadings of each SNP. We carried out this analysis on 38 quantitative traits for which we had GWAS summary statistics that were matched between the European and Japanese Biobanks. After controlling for multiple hypothesis correction, only a single PC (PC10) in a single trait, total bilirubin, was significantly associated with effect size using the BBJ GWAS results, but 24 different PCs across 14 different traits were significantly associated with PCs using the UK Biobank (UKB) GWAS results (<u>Supplementary Table 5</u>, <u>Supplementary Table 6</u>, Extended Data Fig. 8), suggesting that residual stratification remains an issue with using GWAS from the UKB but not from BBJ.

A second measure that we took to increase the portability of BBJ-based associations to the target populations at the expense of statistical power was to limit our analysis to significant associations (GWAS *P*<1×10^−6^), as well as to the sign of the effect on the trait alone. Across a large number of matched traits between BBJ and UKB^84,85^, we found that >95% of all significantly-associated alleles have the same direction of effect. In contrast, effect sizes between BBJ and the UKB were only correlated at ∼70% (<u>Supplementary Table 7</u>), and so effect sizes appear to be less portable. Second, evidence from model organisms, particularly from plants where over 300 studies have been conducted with isogenic lines grown across different environmental conditions, suggest that across a range of traits, while the effect size of QTLs vary, effect directions are almost entirely conserved (98% consistency in effect direction across all comparisons tested)^86^.

In summary, we tested for polygenic selection in a way that reduces statistical power but is more robust to confounding. We identify a set of variants significantly associated with a trait in a Japanese sample, whose population structure and thus potential for population stratification is uncorrelated to that in our ancient European sample. We then carried out a test for selection by comparing the chi-squared statistic of trait-associated alleles, considered alongside with the direction of change in frequency, to those of random SNPs (<u>Methods</u>).

### Signals of polygenic selection that differ across epochs of European history

In each of the three epochs, we tested for selection by comparing our selection statistic in trait-associated SNPs to a distribution of matched controls resampled 10,000 times. The control was composed of an equal number of SNPs matched for deciles of derived allele frequency, recombination rate^87,^ and intensity of background selection^88^ (<u>Methods</u>). We restricted the 220 total traits in the Biobank of Japan to only those that had at least 20 SNPs significantly (*P*<1×10^−6^) associated with the trait. To assess the directionality of genetic change, we polarized our selection statistic to the direction of the effect allele in the GWAS (polarized chi-squared statistic) and asked whether the mean observed polarized statistic for a trait was below the 2.5 or above the 97.5 percentiles of all the matched null samples. In total, we identified 39 traits that reach this level of significance across the three epochs (Fig. 4, <u>Supplementary Table 8</u>). In carrying out this analysis, we checked that null distributions for all traits were approximately normally distributed (Extended Data Fig. 10), and that we had enough variants to prevent the same variant being sampled multiple times (<u>Supplementary Table 9</u>).

**Fig. 4:**
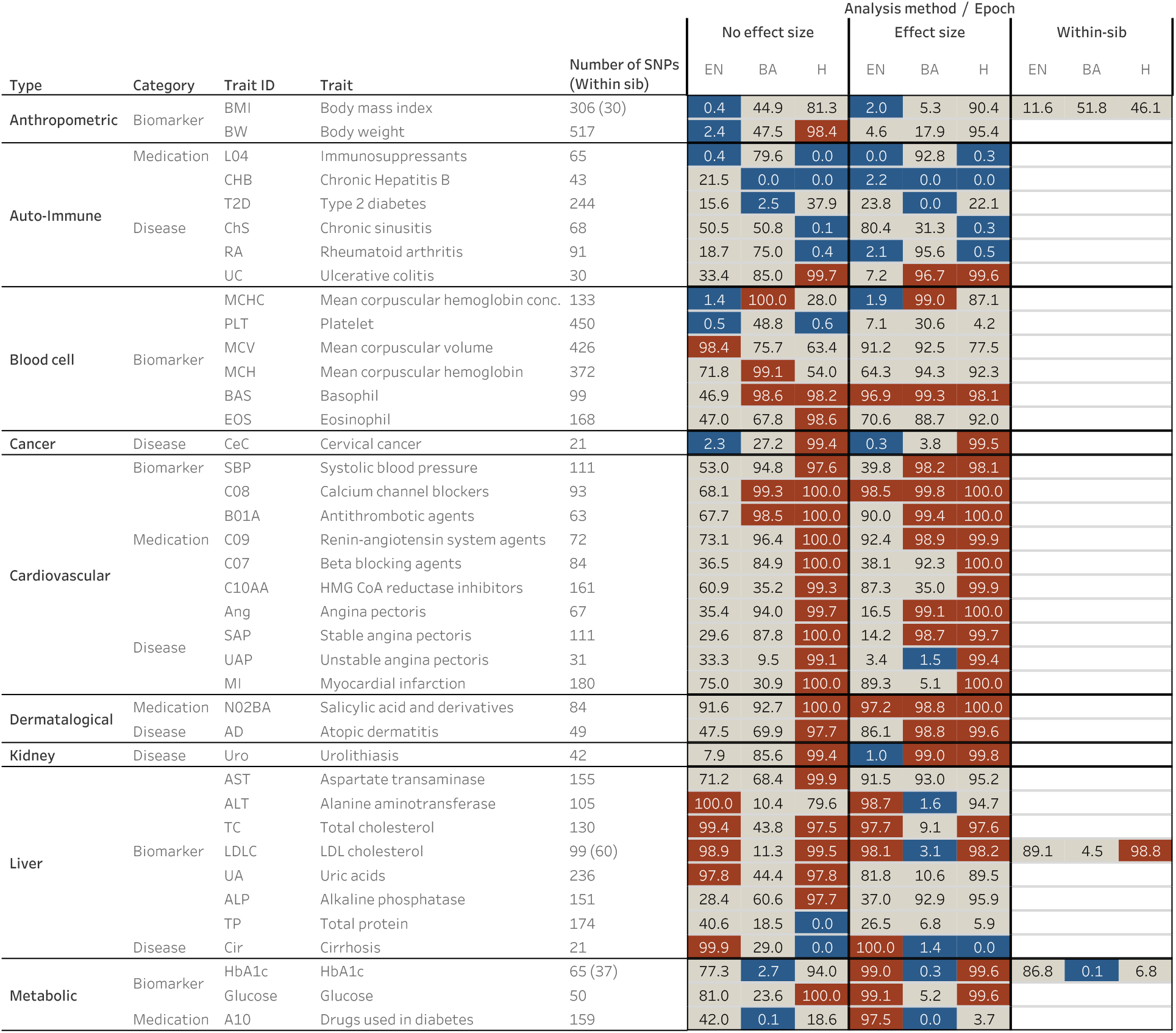
Signals of polygenic selection. Traits shown in red in a given epoch are ones for which we find evidence for post-admixture selection favoring trait-increasing alleles during that epoch. Traits shown in blue show evidence for trait-decreasing in that epoch. In gray are non-significant results. The within-sib GWAS results are only available for a small subset of traits that overlap with BBJ and have a greatly reduced number of SNPs that are genome-wide significant in the GWAS (SNP counts shown in brackets where available). These rows with unavailability of GWAS results for traits from within-sib GWAS are left blank.

In the hunter-gatherer to farming transition, we detected evidence for selection favoring body mass-decreasing and cholesterol-increasing alleles. We also found evidence for selection on traits related to blood cell biomarkers such as platelet and hemoglobin concentration.

In the Bronze Age, we detected evidence for selection on alleles associated with some disease endpoints such as hepatitis and ulcerative colitis, as well as several blood and blood-pressure-related traits. We also observed evidence for selection favoring alleles increasing triglyceride levels.

The vast majority of polygenic adaptation signals we observe were in the Historical period, though several of the traits we identify could be genetically correlated. Importantly, we detect selection favoring alleles that increase rates of heart disease due to myocardial infarction (heart attacks). In addition, we detected evidence for selection on alleles associated with related phenotypes like angina, as well as biomarkers for cardiovascular disease and cardiovascular prescriptions such as beta-blockers. Finally, we observed signals of selection favoring alleles that increase risk for several common auto-immune diseases.

To investigate alternative ways of carrying out these analyses, we repeated these studies by including effect sizes in the polarized chi-squared statistic; we found that the majority of our signals were replicated using the magnitude of effect as well as direction (Fig. 4, <u>Supplementary Table 10</u>). The cases of non-replication were largely sub-significant (for example, ∼2% vs ∼5% for body weight). We also carried out an additional sensitivity analysis by carrying out the same scan, but this time removing SNPs that were in the lowest 27.5% of the chi-squared statistic distribution (Extended Data Fig. 11). These are positions where the direction of frequency change may have been mis-estimated. Therefore, by restricting to sites that are deviating more significantly from expectation, we might increase our confidence in our estimates of the effect direction but reduce power as the number of SNP positions that we could use in the analysis would decrease. This analysis replicated 90% of the original signals across all epochs (<u>Supplementary Table 11</u>). Importantly, we did not detect evidence for natural selection on height, perhaps because of lack of power or because previous analyses may have been confounded by population stratification^4,89^. A final sensitivity analysis we carried out was to repeat the polygenic selection analysis using summary statistics that were identified from a large consortia study of 25 phenotypes carried out using a within-sibling GWAS design^90^. In theory, family-based GWAS designs can control for demographic and indirect genetic effects, but even with the relatively large number of samples, only 6 out the 25 traits had more than 20 SNPs that met our inclusion *P* value threshold. Out of these 6, only 3 traits overlapped with traits that were also seen in the Biobank of Japan dataset. Largely, the within-sib GWAS data agreed with our previous analysis, with the exception of selection favoring BMI-decreasing alleles in the Neolithic. However, the within-sib analysis was considerably underpowered compared with using the BBJ dataset, as the total number of SNPs were different by an order of magnitude (<u>Supplementary Table 13</u>).

## Discussion

Our analysis highlights the power of ancient DNA time series data to reveal evidence of natural selection in humans that has later become obscured by subsequent evolution. To evaluate the extent to which our results replicate previous findings, we compared our candidate targets of selection with two previous studies. The first, Mathieson et al.^4^, used ancient DNA and a similar approach to ours—detecting sites with unusual allele frequencies compared to genome-wide admixture proportions—but used modern samples from the 1000 Genomes Project as a target population^4^. The second, Pybus et al.^91^, used a composite approach integrating multiple classical selection tests on modern European genomes^91^. We found only one candidate shared between our study and with modern genomes from the ancient DNA based scan in the EN (the HLA region) and two in the BA period (the HLA region and *SLC45A2*), but 9/12 of Mathieson et al.’s^4^ candidates were shared with our Historical epoch candidates (Fig. 3). Similarly, Pybus et al.^91^, found none of the candidates we found in the EN epoch, one in the BA epoch, but 7 that match our candidates from the Historical period. A possible explanation for this is that the admixture in Europe over the past 10,000 years has obscured signals of selection that occurred before the immediate past^14^.

Our approach looking at time trajectories of alleles over a 10,000-year old period also made it possible to assess the impact of alleles in the germline over long time scales. As an example of this, we studied the frequency trajectory of the *CCR5-Δ32* variant, which in homozygous form provides protection against HIV in European individuals. We studied the frequency of this allele using a proxy SNP (rs73833033) that is in high LD with it. Across the 3 epochs, we find no evidence for selection of this allele (p=0.55, 0.05, 0.34 for the EN, BA, and H epochs respectively) in line with the evidence from modern samples, despite previous reports of selection at this locus^92,93^ (Extended Data Fig. 4).

It is important to recognize that the number of candidates we find in each epoch does not reflect the intensity of natural selection in that time. Rather, many factors feed in to epoch-specific statistical power. Consider the example of *LCT*: it is possible that 6,000-3,000 years ago, selective pressures on lactase persistence have been stronger than in the Historical period. Here, selection overcame genetic drift and drove the very rare allele to a frequency of several percentage points of the population. Yet, the largest change in allele frequency, from a few percent to the majority allele in northern Europe, only occurred in the Historical period. These are the changes that we are most powered to detect. Another important caveat is that the genomic control and null model we rely on may not be equally adequate in all parts of the genome, particularly in the HLA region where mutation rates, recombination rates, natural selection, and genetic drift are highly atypical^94^. Nevertheless, our estimation of genome-wide admixture proportions using *qpAdm* suggests that our expected frequencies broadly capture the allele frequency shifts associated with admixture.

Our results also allow us to interpret our signals of selection in light of archaeological, evolutionary, and biological evidence. In particular, they allow us to test theories about gene-culture co-evolution, specifically with regard to hypotheses about how major changes in lifestyle in the last ten millennia in Europe have or have not resulted in signals of genetic adaptations.

One important set of insights relates to the genomic impact of the major transition from hunting and gathering to farming. A set of alleles that were targets of selection in this period have to do with decreased body weight/size. Complementarily, the archeological record also shows an overall decrease in body size during the Neolithic transition^95^. One hypothesis is that a reduction in overall calorie intake, a trait that would be genetically correlated with reduced body weight, was advantageous in the Neolithic when famines and resource instability became more frequent^96^. Similarly, selection for more efficient storage and use of glucose in tissues during periods of famine or pathogen outbreaks might also underlie several of our selection signals associated with insulin secretion, regulating glucose in the blood stream.

Our results also allow us to re-examine the hypothesis for selection on the lactase persistence allele. A recent study suggested that milk exploitation and consumption started well before the lactase persistence allele began to be selected, and that the selected allele did not show consistent associations with improved fitness or health in modern individuals, perhaps suggesting that the ability to digest lactose into adulthood was only selected for in conditions of food scarcity^97^. While this study was exclusively restricted to just this allele and phenotype, our genome-wide scan adds additional perspective on the rationale for selection at this locus by connecting it to selection at other candidates in the same time period. In the Historical period, along with the *LCT* locus, we detect selection candidates in *SH2B3* and *DHCR7*—two genes that are directly related to vitamin D binding as well as a candidate in *SLC45A2*, a major locus of light skin pigmentation in Europeans, a phenotype which also promotes vitamin D synthesis from sunlight. Taking these results together, our results may suggest an alternative to the caloric supplementation hypothesis; namely, that selection acted to increase calcium uptake—via improved vitamin D absorption as well as increased dietary uptake through the consumption of milk—a finding that has also been discussed in another recent study^13^. Vitamin D is almost entirely absent in a plant-based diets, and these results might also help explain the differences in both lactase persistence and skin pigmentation between Northern and Southern Europe, with increased sun exposure in Southern Europe allowing for sufficient vitamin D synthesis despite a similar dietary transition.

A third major set of loci we find as candidates are involved in pathogen response or are expressed on the cell surface of immune cells. A hypothesis behind selection at these loci could be related to the potentially increased infectious disease load in the Neolithic brought about from people living in closer proximity to animals as well as to each other, a set of pressures that would have become dramatically stronger as population sizes increased exponentially in the Bronze Age and Historical periods. Indeed, over the past few years, a number of ancient DNA studies have reported pathogen sequences from the Neolithic period and later^98–102^. These studies revealed past epidemics and found evidence for adaptation of viruses and bacteria to the human host. Evidence from population history analysis also shows that Europe in the past 10,000 years has seen large scale migrations into the European subcontinent from individuals practicing different lifestyles. Signals of selection in these immune loci could be reflect the arrival of new zoonotic pathogens that arrive with incoming farmers and pastoralists who brought domesticated animals with them (sheep, cattle, and goats in the case of the first farmers^103^, and horses in the case of Steppe Pastoralists^104^). Our findings of pervasive upward shifts in the frequencies of alleles increasing cardiovascular disease and auto-immune disease risk can also be interpreted in this light. All else being equal, our findings suggest that today, people with hunter-gatherer genomes would have been at lower risk for cardiovascular and autoimmune disease. The high prevalence of cardiovascular disease in modern societies could be in part due to past selection for heightened inflammatory response—the immune system’s primary response to harmful stimuli including pathogens^105^. That is, beginning in the Neolithic and intensifying in the subsequent periods, humans were subject to a greater infectious load, and selection for proinflammatory genes and a strong inflammatory function due to the secretion of adipokines, which underlie cardiometabolic diseases, may have resulted in increased risk for cardiovascular disease.

While we find evidence for two hypotheses concerning gene-culture co-evolution in the last ten thousand years—with selection for traits related to metabolism as well as immune response—we did not have power to detect selection for cognitive or neuro-psychiatric disease traits, due to the limited data and relatively small sample size for these traits in the Biobank of Japan data. There is no evidence in the genetic data for selection on such traits, but future larger studies might provide power to detect such signals.

While our work offers some methodological improvements compared to previous efforts, the greatest improvement in resolution comes from the quality and quantity of ancient DNA data. More ancient genomes are becoming available from different geographic regions and time periods. Extending the type of analysis we report here to these datasets has the potential to enrich our understanding of the history of natural selection on humans beyond what could be learned through analyses of contemporary sample alone, where ancient selective events are obscured by admixture and drift, and where their timing cannot be directly determined.

## Methods

### Ancient DNA data curation

We obtained ancient DNA sequencing data from the Allen Ancient DNA Resource (https://reich.hms.harvard.edu/allen-ancient-dna-resource-aadr-downloadable-genotypes-present-day-and-ancient-dna-data, version 51), and subsetted the data to only include samples that were enriched for ∼1.2 million nuclear targets with an in-solution hybridization capture reagent.

To analyze the data, we began with the raw read data for all of these samples and sorted the read pairs by searching for the expected identification indices and barcodes for each library, allowing up to one mismatch from the expected sequence in each case. We removed adapters and merged together sequences requiring a 15 base pair overlap (allowing up to one mismatch), taking the highest quality base in the merged segment to represent the allele. We mapped the resulting sequences to the hg19 human reference genome^106^ using the same command of BWA^107^ (version 0.6.1). We removed duplicate sequences (mapping to the same position in the genome and having the same barcode pair), and merged libraries corresponding to the same sample (merging across samples that the genetic data revealed were from the same individual). For each individual, we restricted to sequences passing filters (not overlapping known insertion/deletion polymorphisms, and having a minimum mapping quality 10), and trimmed two nucleotides from the end of each sequence to reduce deamination artifacts. In addition, we also restricted to sequence data with a minimum base quality of 20.

We assessed evidence for ancient DNA authenticity by measuring the rate of damage in the first nucleotide, flagging individuals as potentially contaminated if they had less than a 3% cytosine-to-thymine substitution rate in the first nucleotide for a UDG-treated library and less than a 10% substitution rate for a non-UDG-treated library. We used contamMix to test for contamination based on polymorphism in mitochondrial DNA^108^ and ANGSD to test for contamination based on polymorphism on the X chromosome in males^109^, removing individuals with evidence for contamination. For population genetic analysis to represent each individual at each SNP position, we randomly selected a single sequence (if at least one was available). For the selection analysis, in order to obtain read count information on a per sample basis, we used BCFtools^110^ version 1.3.1 to obtain reference and alternate counts at each genomic position.

### Principal components analysis

We carried out PCA using the smartpca package of EIGENSOFT 7.2.1^111^. We used default parameters and added two options (lsqproject:YES and numoutlieriter:0) to project the ancient individuals onto the PCA space. We used 991 present-day West Eurasians^22,112,113^ as a basis for projection of the ancient individuals. We also computed F_ST_ between groups using the parameters inbreed:YES and fstonly:YES. We restricted these analyses to the dataset obtained by merging our ancient DNA data with the modern DNA data on the Human Origins Array and restricted it to 597,573 SNPs. We treated positions where we did not have sequence data as missing genotypes. Fig. 1 shows the PCA of all ancient samples. Extended Data Fig. 1 shows underlying modern samples used for the projection, along with the ancient individuals.

**Extended Data Fig. 1.**
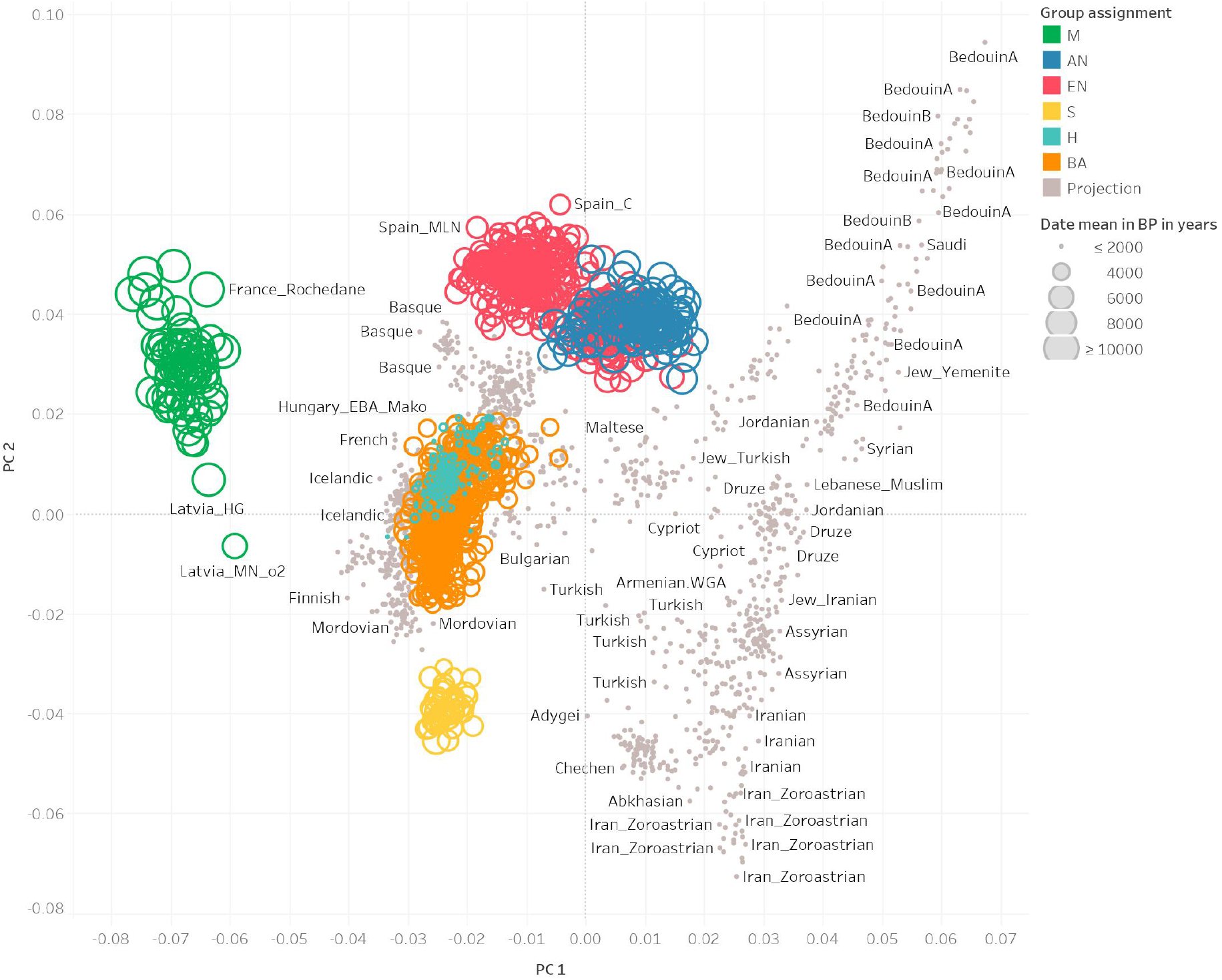
PCA of ancient samples as well as the basis set of modern samples used in the projection analysis (in grey).

### Population grouping and f-statistics

We grouped samples into several broad genetic and cultural categories that represented the major ancestry groups observed in our European time transect. Our group assignments were:

M, Mesolithic - Hunter-Gatherers with no evidence of any admixture from Anatolian Farmers dated to an average of around 8,600 BP with the majority from Eastern Europe

AN, Anatolia Neolithic - Farmer samples dated to an average of around 7,400 BP largely from present day Turkey, Greece, and the Balkans, with little to no European Hunter-Gatherer admixture

EN, Europe Neolithic - Farmer samples dated to an average of around 5,400 BP with the majority of samples from Central and Western Europe

BA, Bronze Age - Samples dated to an average of around 4,000 BP largely from the Bell Beaker cultures of Czech Republic, Great Britain, Germany, and Slovakia

S - Steppe Pastoralists dated to an average of around 4,800 BP with many from Yamnaya and Afanasievo cultures of the Eastern Steppe

H, Historical era - Samples dated to an average of around 2,000 BP with the vast majority from England and Scotland, as well as a minority of samples from Central Europe

### Admixture modeling of ancient Europeans

We used *qpAdm* from ADMIXTOOLS to estimate the mixing proportions for the ancestral populations of each model^113^. *qpAdm* estimates the mixing proportions using the expected values of *f*_4-_ statistics, where *f*_4_(*A, B*; *C, D*) represents the correlation in allele frequency differences between the groups (*A, B*) and (*C, D*)^114^. We used seven outgroups for the computation of the *f*_4_-statistics: Ethiopia_4500BP, Russia_Ust_Ishim_HG_published.DG, Russia_MA1_HG.SG, Israel_Natufian, Italy_North_Villabruna, Iran_Ganj_Dareh_N, and Russia_Boisman_MN.

We leveraged previous work that provides a demographic model for major ancestry transitions in Europe^4,22,23,27^. We modeled European Farmers (EN) as a 84% mixture of early farmers from Anatolia (AN) and 16% mixture of European Hunter-Gatherers (M). We modeled European Bronze Age samples as a 48% mixture of European Farmers (EN) and 52% mixture of Steppe Pastoralists (S). Finally, we modeled Historical era samples from Europe (H) as a 85% mixture of Bronze Age samples and a 15% mixture of European Neolithic samples, reflecting the additional ancestry changes at that time.

### Genome-wide scan for natural selection

To estimate the population allele frequencies at each site, we obtained the maximum likelihood estimate from the likelihood of a given frequency *p* using an approach first described in Mathieson et al. 2015. Let *p* be a reference allele frequency, *R*_*i*_ be the number of reads with the reference allele, *T*_*i*_ be the total number of sequences, *N* be the number of samples for the population, and ε be a probability of error. Let the binomial probability mass function be denoted as 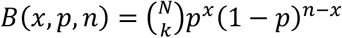. Then the likelihood of a frequency *p* given the read data is

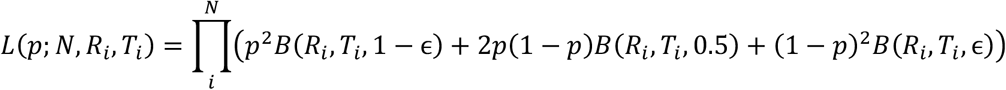

Minimizing the negative log-likelihood function produced the allele frequency estimate for each population at every site. Samples with 0 reference and alternate reads at a site were excluded from the calculation of the maximum likelihood estimate. We used the SLSQP solver from SciPy^115^ to minimize the negative log-likelihood function, setting the bounds for the allele frequency at 0.01 and 0.99. We also removed all positions where all reads were missing in any of the populations used in the scan.

The expected frequency of the target population was also obtained given the mixing proportions and estimated frequencies of the ancestral populations. For instance, suppose we have the admixture model C = αA + (1 − α)B. Then under neutrality, the expected allele frequency for population *C* would be *p*_*E*_ = α*p*_*A*_ + (1 − α)*p*_*B*_, where *p*_*A*_ and *p*_*B*_ are the observed allele frequencies of populations *A* and *B*, respectively.

Let *p*_*E*_ be the expected frequency of the target population computed as the sum of the products of the allele frequencies of the ancestral populations and their mixing proportions, and let *p*_*O*_ be the observed frequency of the target population. We tested when the observed allele frequency deviated from expectation using the likelihood ratio test.

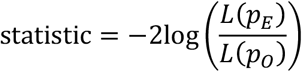

This statistic was used to compute a *P* value from the 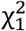 distribution. To address genomic inflation, a control factor was applied to the statistics such that 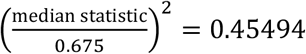 after removing 49,000 SNPs of functional importance^4,31^. We also removed genomic positions that were covered by >15,000 reads (coverage >10x mean coverage) across our dataset, due to potential mis-mapping artifacts.

Previous work has examined the robustness of this particular approach to mis-estimation of allele frequencies as well as sample sizes, but we added additional power calculations to our particular scenario. First, we carried out an analysis where we modified the ancestry proportions in 5% increments from the actual proportion and again examined the number of 1Mb regions that remained significant according to our criterion after genomic inflation correction. Our results suggest that we are well-powered to detect the majority of our signals even with mis-specification of the admixture proportion by over 30% (Extended Data Fig. 2).

Second, we carried out a sub-sampling analysis where we down sampled the overall dataset in 5% increments (that is, reducing the sample sizes of both the two source populations and the target population across all 3 epochs in steps of 5%), and then examined the number of 1 Mb regions that remained significant according to our criterion after genomic inflation correction. We see that with 90% of the data, we are essentially recapturing most of our signals, though the lack of a clear plateau in our analysis suggests that increasing sample sizes further is likely to continue to improve the power to discover new loci (Extended Data Fig. 3). Small increases in power are seen even at slightly larger sample sizes, as our sampling process is carried out at the level of individuals. This is as expected, as coverage varies greatly by sample and across genomic position, but the overall upward trajectory of increased power with increased sample size is clear.

**Extended Data Fig. 2.**
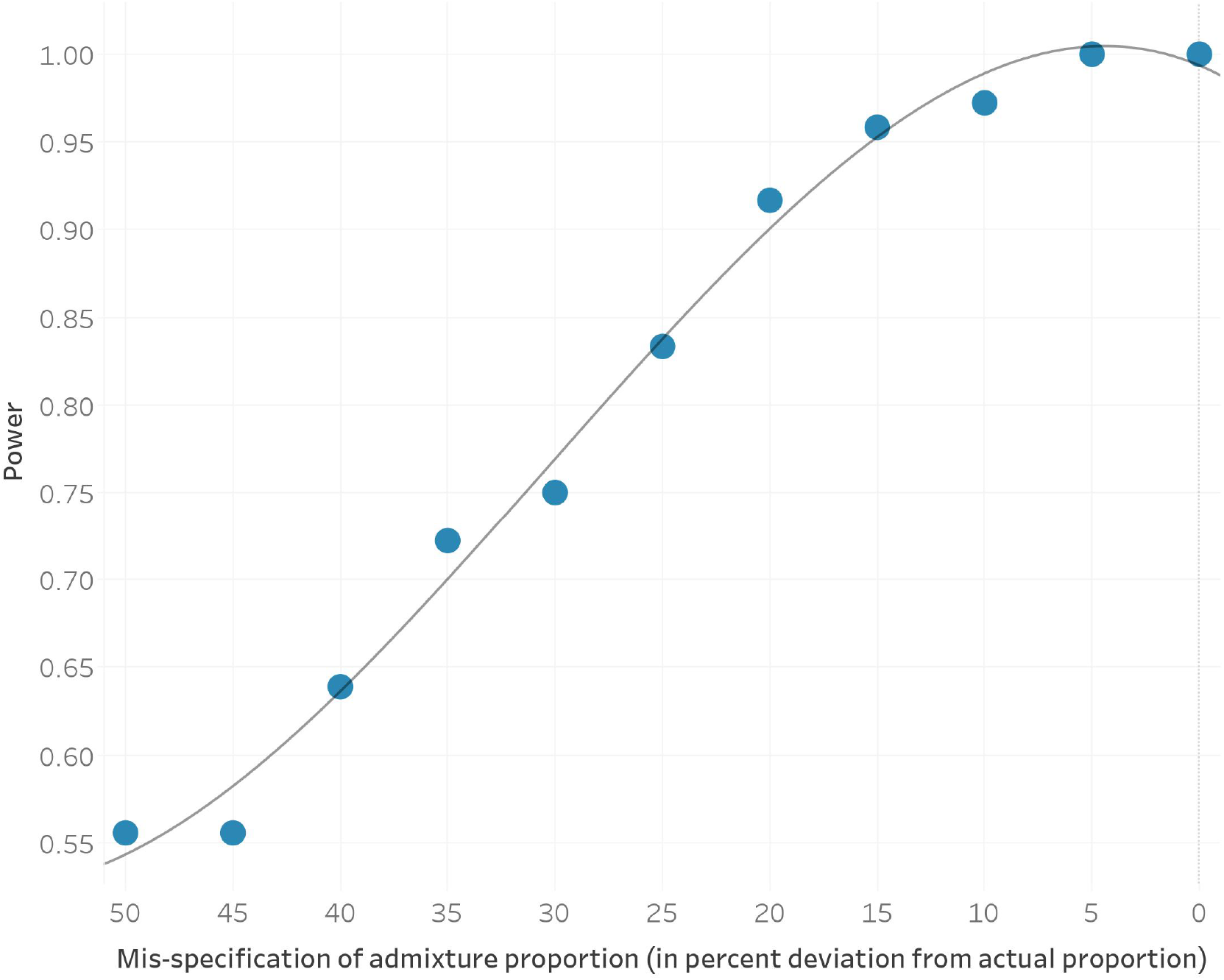
The power to discover significant genomic regions as a function of admixture proportion mis-specification shown in percent deviation from the actual mixture proportion. Grey line shows the quadratic fitted estimate.

**Extended Data Fig. 3.**
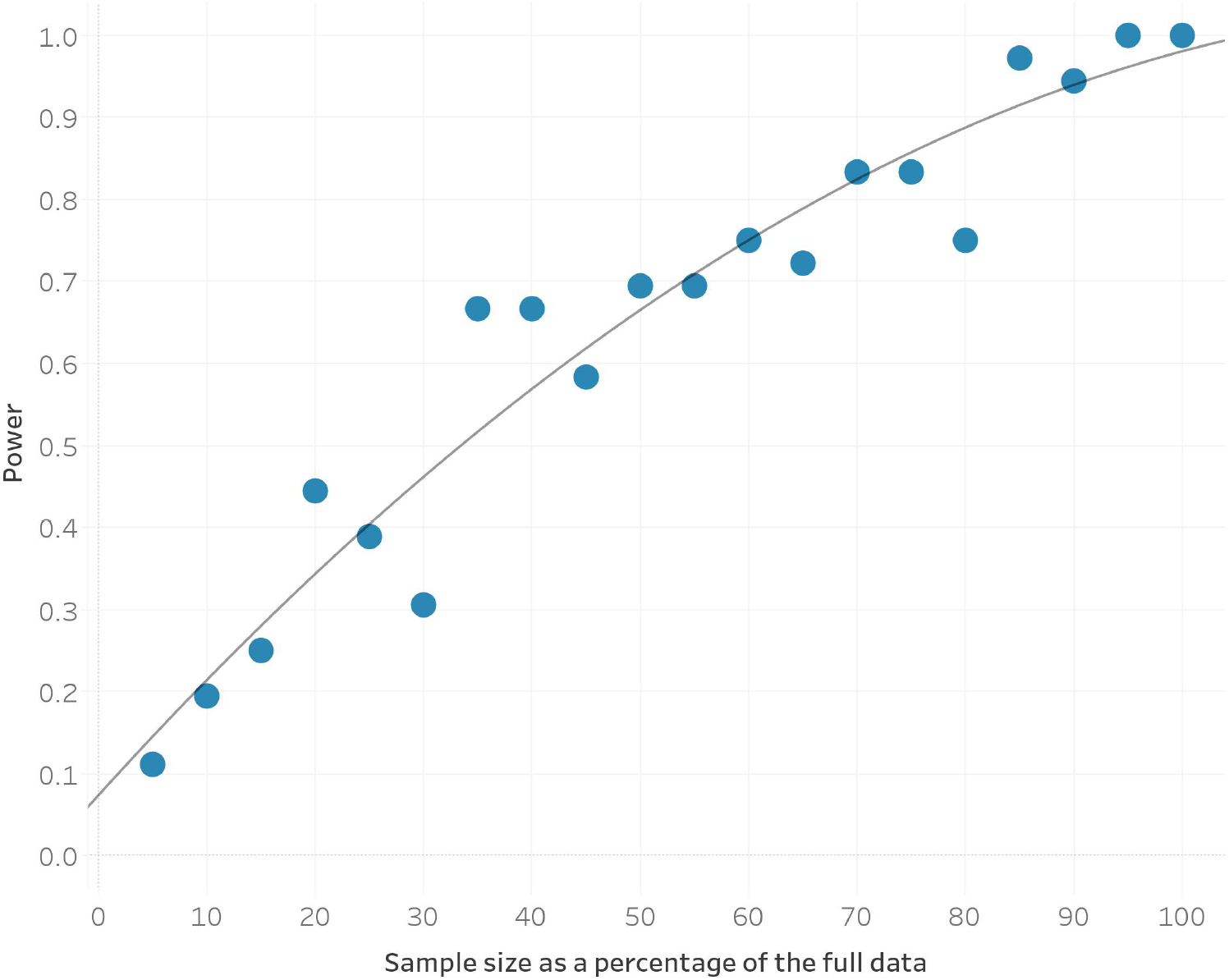
The power to discover significant genomic regions as a function of sample size (here reported as a percentage of the full dataset of 1,291 samples). Grey line shows the quadratic fitted estimate.

**Extended Data Fig. 4.**
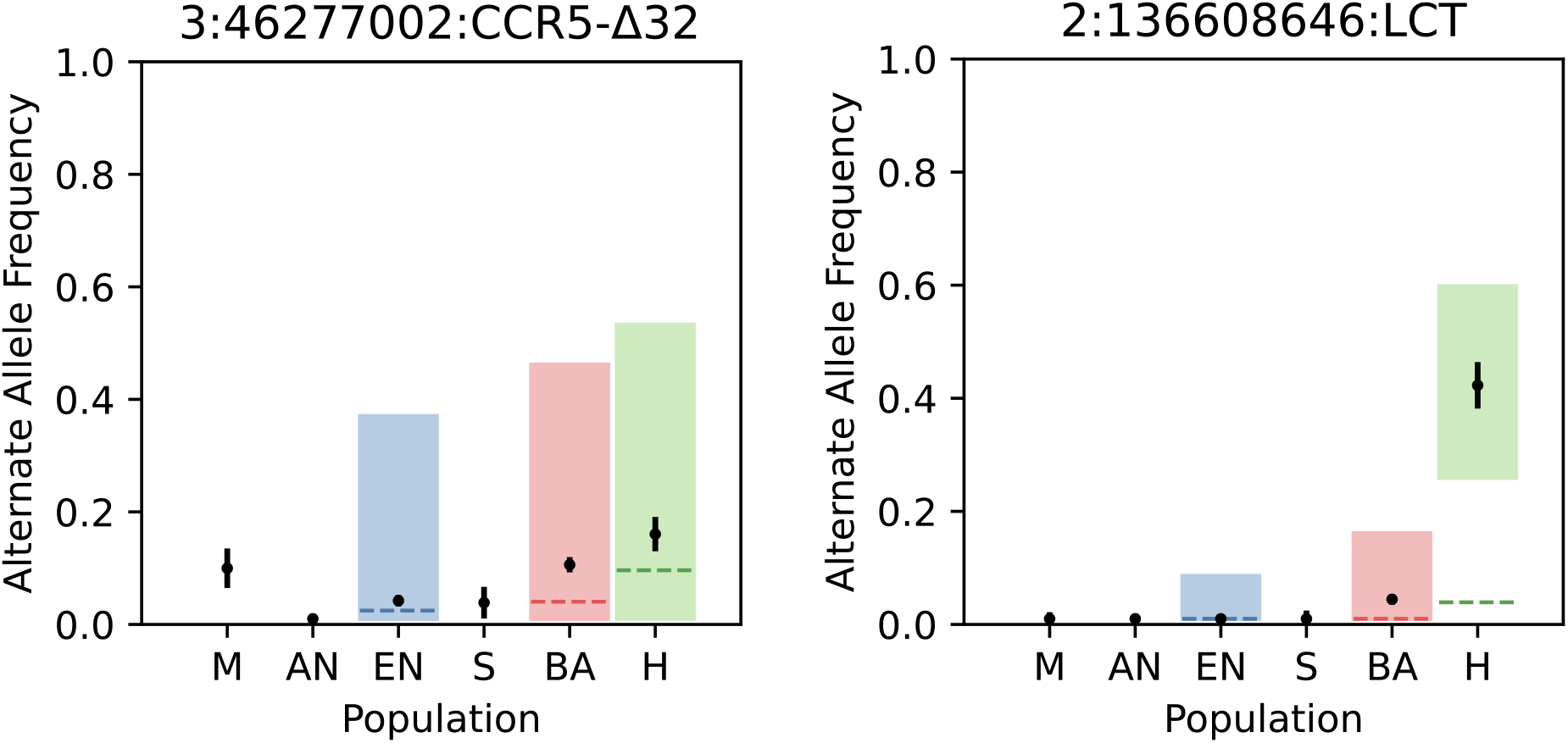
Point estimates and standard errors of alternate allele frequencies in each population. The dashed lines (blue for EN, red for BA, and green for the H epoch) are the expected allele frequencies of the alternate allele based on genome-wide expectations of admixture proportions. An expected allele frequency that falls outside of the shaded regions would result in a significant *P* value from the likelihood ratio test after correction for genomic inflation. The *CCR5-Δ32* allele does not appear to be under selection in any of the epochs, but the *LCT* allele shows major changes in frequency in the Historical period.

**Extended Data Fig. 5.**
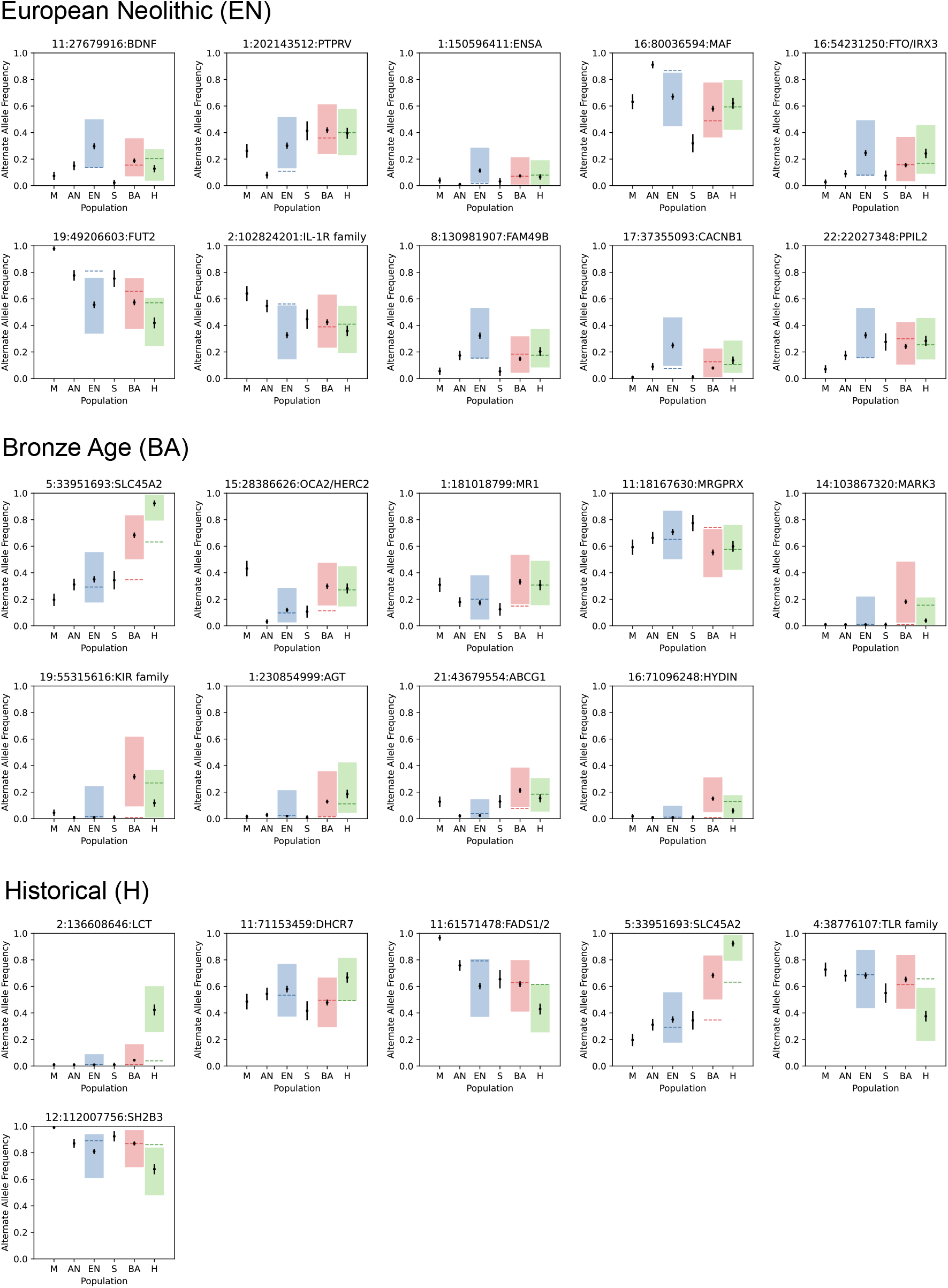
Allele frequency and 95% confidence intervals for selected variants across the 3 epochs. The dashed lines (blue for EN, red for BA, and green for the H epoch) are the expected allele frequencies of the reference allele based on genome-wide expectations of admixture proportions. An expected allele frequency that falls outside of the shaded regions would result in a significant *P* value from the likelihood ratio test after correction for genomic inflation.

### Correlation of ancient selective events to those seen in modern Europeans

Selective events that occurred further back in time might be obscured by drift, admixture, or fluctuations in selective direction over the generations. We wanted to examine if the signals we uncovered through our ancient selective scan might also have been seen in selection scans of modern Europeans. In order to do this, we compared the chi-squared statistic we obtained at every locus across all epochs to a machine learning-based ensemble classifier that integrates several different classical selection tests into a single predictor^91^ in two ways. First, we computed simple overlaps of regions under selection using the XGBoost algorithm and regions found to be under selection in our analysis. Second, we examined the number of loci that overlapped with a previous ancient DNA based scan for natural selection^4^.

### Variant effect predictor

For each population, we filtered the SNPs to a list of variants that had corrected *P* values above a genome-wide significance level of 5 × 10^−8^ and at least two other SNPs above the significance cutoff within 1 Mb. We then used the Ensembl Variant Effect Predictor^116^ to obtain a list of the nearest genes associated to each variant and filtered to retain only protein coding genes. All annotations, frequencies and enrichment analysis were performed on the human reference genome build GRCh37^116^. We include significant loci in our selection scan and genomics annotations in a 1 Mb neighborhood using LocusZoom^117^ (Extended Data Fig. 6).

**Extended Data Fig. 6.**
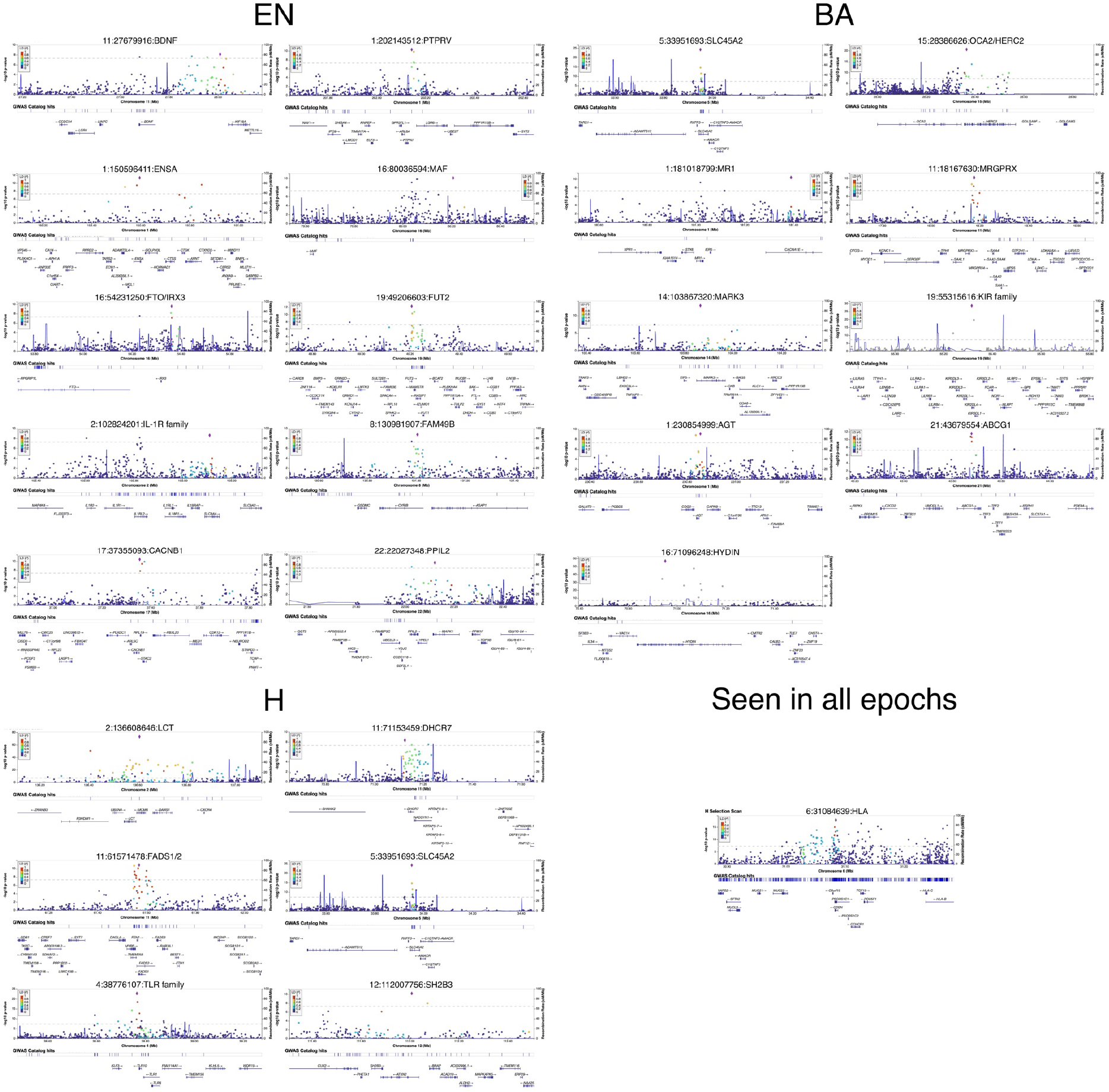
LocusZoom plots of all selected variants and gene annotations in a 1 Mb region around them.

### Enrichment analysis

We used the Functional Mapping and Annotation of Genome-Wide Association Studies tool to obtain significant gene sets for each epoch. The gene sets were produced by comparing the genes of interest against sets of genes from MsigDB using hypergeometric tests. We performed this analysis for gene sets from the GWAS and GO functional categories^34^.

### Polygenic selection

We used a modified version of the test from Choin et al.^118^ to test for evidence of polygenic selection. We used GWAS summary statistics from the UK Biobank to test for selection in a control set of traits, which included skin color, hair color, and triglycerides^119^. We used summary statistics from the Biobank of Japan to test for selection in 220 different traits^120^. For each trait, we classified each allele as trait-increasing or trait-decreasing using the effect direction. We then polarized our admixture scan selection statistic such that a positive sign indicated directional selection of the trait-increasing allele. In other words, for a given loci *i*, our polarized statistic was computed as | *χ*_*i*_ | sgn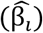 where | *χ*_*i*_ | is the magnitude of the chi-squared statistic from the monogenic selection scan, and sgn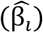 is the sign of the effect size for the allele that increased or decreased from expectation. For each trait, we compared the mean polarized statistic of the GWAS significant SNPs (at significance level *P*<1×10^−6^) to the distribution of the mean polarized statistics of randomly sampled SNPs (Extended Data Fig. 7). The rationale for this test is that trait-associated SNPs would be more likely than random to undergo short-term directional selection.

**Extended Data Fig. 7.**
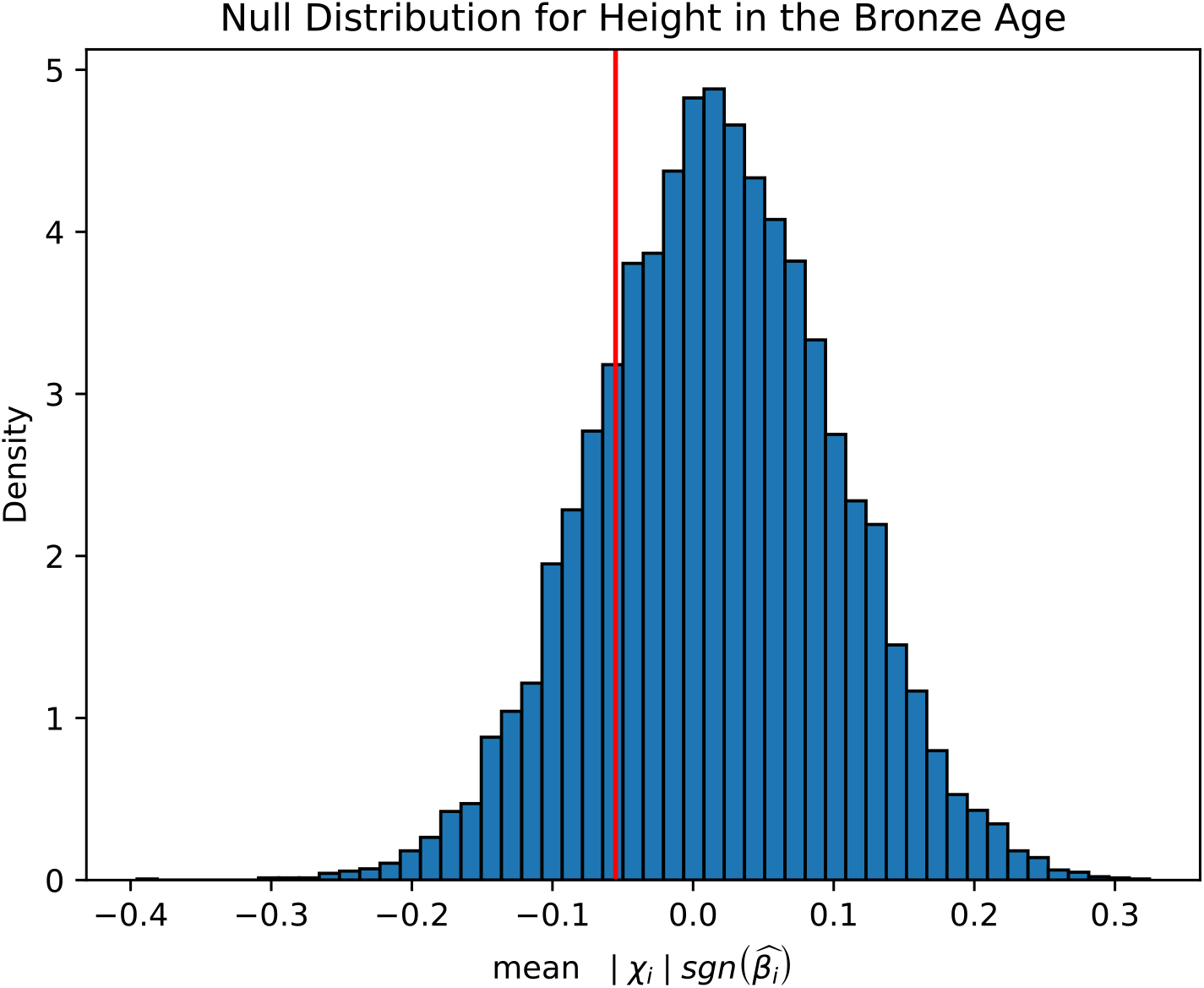
Null distribution of random subsamples of matched controls (in blue bars) and the observed statistic (the red line) for a single trait, height, in the Bronze Age. Observed statistics below the 2.5th percentile or above the 97.5th percentile of the null were considered significant.

To investigate the impact of population stratification on the GWAS effect sizes, we regressed GWAS effect sizes on PC loadings on both the UK Biobank (UKB) and Biobank of Japan (BBJ) datasets. In Extended Data Fig. 8, we show the results of this regression on the best studied and most heritable of the traits we examined: height. Our results show that effect sizes from the UKB are significantly associated with PC loadings, but effect sizes from BBJ are relatively uncorrelated with PC loadings, which is in agreement with work from Chen et al^83^. We also observed these results on a set of 38 other matched quantitative traits and found that only 1 trait has a single PC (PC100) that had PC loadings significantly associated with effect size using the BBJ dataset, but using the European GWAS 24, PC loadings across 14 traits were significantly associated with effect size (<u>Supplementary Table 5</u>, <u>Supplementary Table 6</u>).

**Extended Data Fig. 8.**
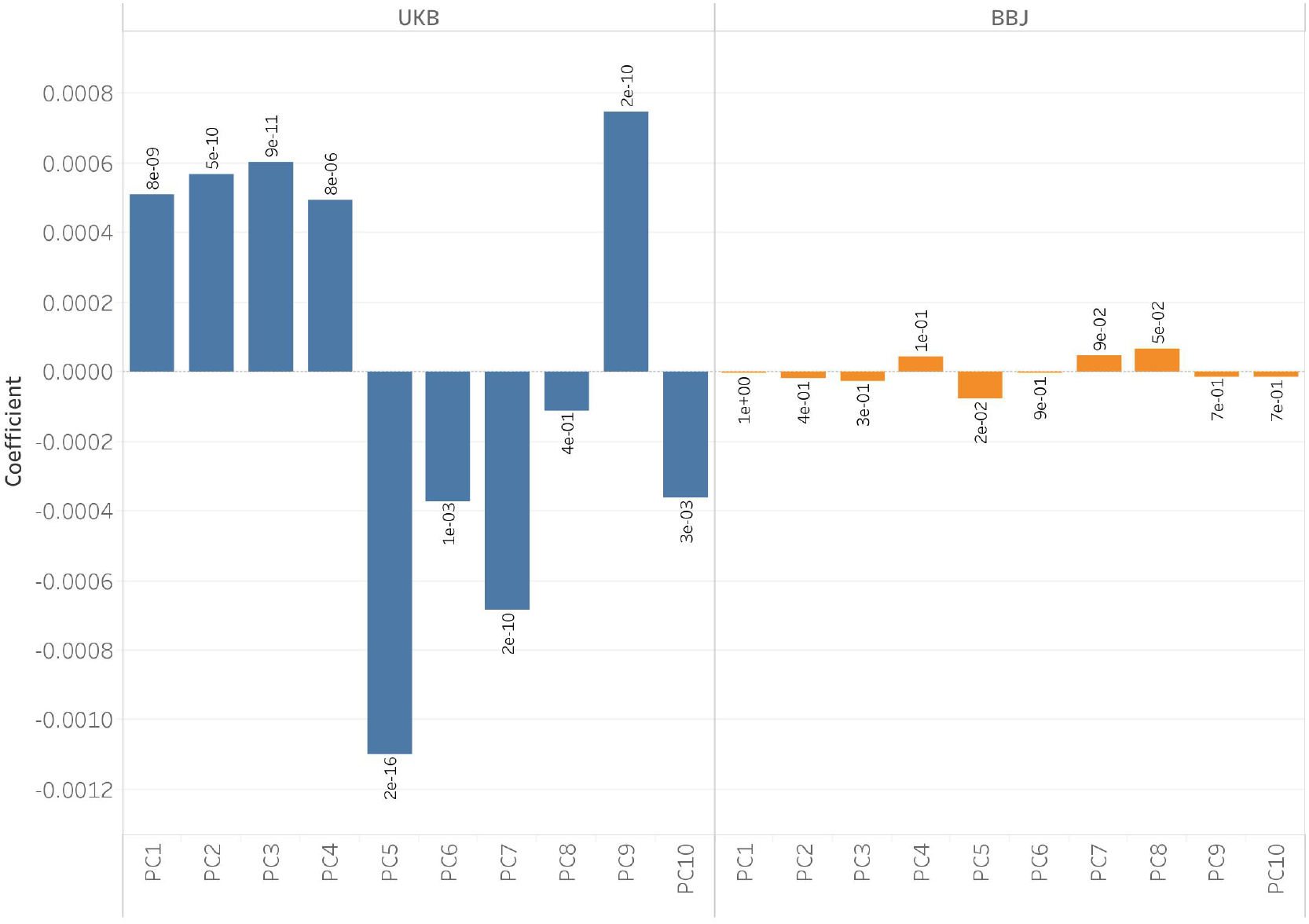
Regression coefficients of the GWAS effect sizes for height from the UKB and BBJ on PC loadings from the West Eurasian basis space generated from diverse samples of modern European genomes. There are no PCs that are significantly associated with effect sizes using BBJ, but this is not the case for UKB where several PCs are significantly associated with the trait.

For each trait, we took 100 kb windows and chose only a single SNP with the lowest *P* value to represent blocks of independent associations with the trait, and we computed the mean polarized statistic of the set of SNPs that were also below a genome-wide significance threshold of 1 × 10^−6^. To match these observed variants to controls, we binned the other variants in the genome based on derived allele frequency, B statistic, and recombination rate. The derived allele frequency bins were separated into 8 equally sized bins that ranged from 0 to 1. The B statistic bins were divided by deciles^121^. The recombination rate bin thresholds were computed such that the recombination rates of the SNPs used in the admixture scans were evenly distributed across 8 bins. In carrying out this empirical sampling procedure, we ensured that matched control distributions were normally distributed (Extended Data Fig. 9), and that this was the case across different bins (Extended Data Fig. 10). To ensure that we very rarely resampled the same alleles each time in our random distributions, we ensured that there were a minimum of 100 variants in each bin (<u>Supplementary Table 9</u>).

**Extended Data Fig. 9.**
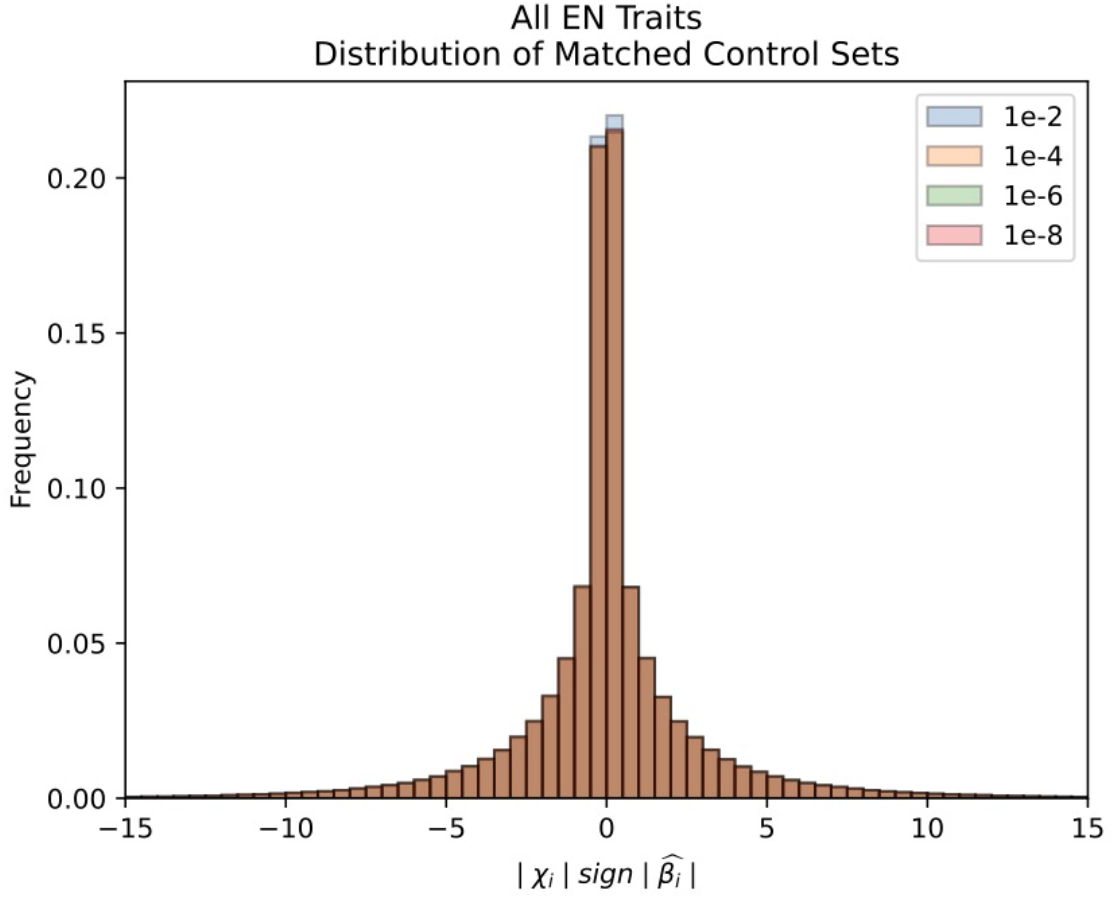
Null distributions across all the traits we tested, across different *P* value thresholds of the GWAS, showed normality in the null distributions and were centered around 0.

**Extended Data Fig. 10.**
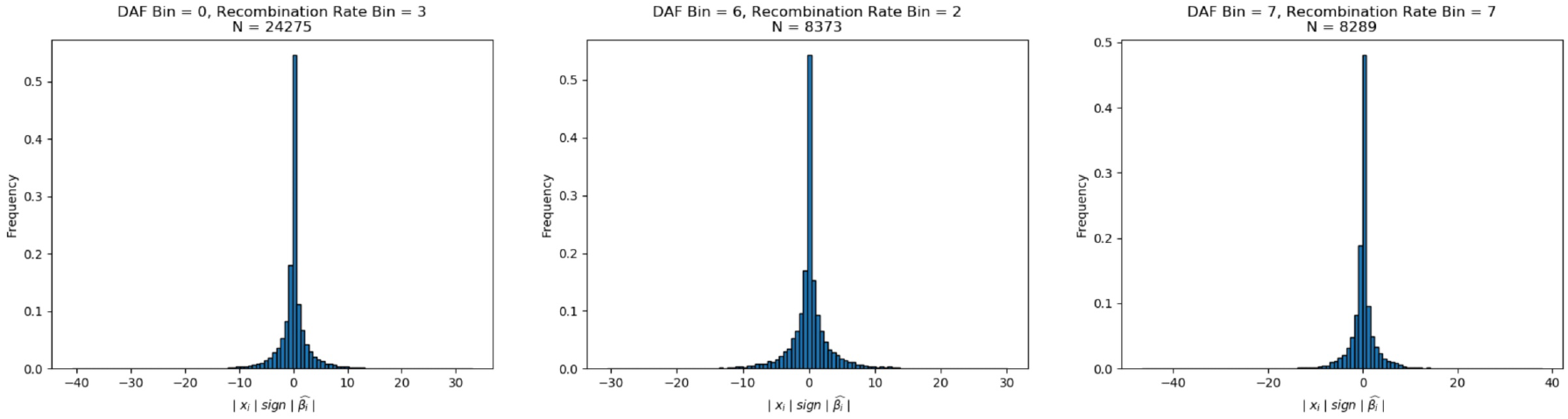
Distribution of chi-squared statistics across different bins also show null distributions centered around 0.

We then randomly sampled variants that were not associated with the trait 10,000 times to match the profiles of the variants with the lowest association *P* values and computed the mean polarized selection statistic for the sampled variants. We considered there to be directional selection in the trait-increasing direction if less than 2.5% of the sampling trials had a mean polarized selection statistic higher than that of the variants with the lowest *P* values. Similarly, we considered there to be directional selection in the trait-decreasing direction if less than 2.5% of the sampling trials had a mean polarized selection statistic lower than that of the variants with the lowest *P* values. We only report significant association on traits that had at least 20 SNPs that were significant at a GWAS *P* value threshold of less than 1×10^−6^).

We also repeated this analysis by multiplying each variant’s chi-squared statistic by its effect size, thereby computing a score that also includes the magnitude of its effect on the trait beyond just looking at its direction. We report all results for this analysis in <u>Supplementary Table 10</u>. Finally, we carried out an analysis reducing the SNPs used by removing sites where the chi-squared value was in the bottom 27.5% of each epoch. We chose to use 27.5% as it was just past the modal value of the distribution at 25%. The positions that were removed by this process are ones where we have a higher likelihood of mis-estimating the direction of frequency change. This analysis replicated the majority (90%) of the original signals. Traits that were not replicated were due to reduction in the overall number of SNPs being reduced to below 20, a condition we required for the sampling process to be reasonable. Only 4 new traits that were not significant previously were seen to be significant, but these were sub-significant (5%) in the original analysis (<u>Supplementary Table 11</u>).

**Extended Data Fig. 11.**
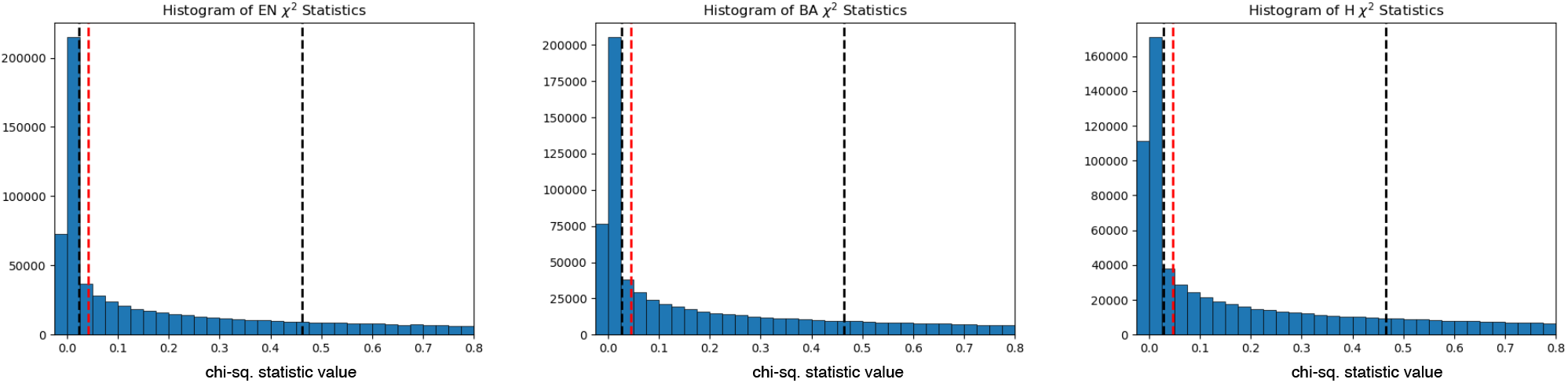
Distribution of the frequency (y-axis) of chi-squared statistic values (x-axis) across epochs. Black lines show the 25th and 50th percentiles and the red line is the 27.5th percentile.

In addition to carrying out analysis with the Biobank of Japan dataset, we repeated the analysis with the UK Biobank dataset <u>Supplementary Table 12</u>. We caution that the results of this analysis are not readily interpretable or comparable with those from the Biobank of Japan in light of the issues we discuss with the applications of GWAS with known population stratification artifacts to detect polygenic selection, but we provide these results for completeness. Finally, we reran the polygenic selection analysis using data from the within-sibling GWAS consortium. The GWAS estimates from within-sibling studies are, in theory, better controlled for issues associated with stratification, but the total number of genome-wide significant loci that met our threshold limited this approach to only a handful of traits. We report the results of this in <u>Supplementary Table 13</u>.

## Code Availability

The code used to run the selection scans for individual alleles and polygenic traits is available at https://github.com/Narasimhan-Lab/1000-genomes-natural-selection.

## Acknowledgements

We would like to thank Robert Maier, Alex Okomoto, Iain Mathieson, and Dan Lieberman for useful discussions. Megan Le was supported by an Undergraduate Research Fellowship Award, an Advanced Summer Research Fellowship Award, and Endowed Presidential Scholarship Awards. Olivia Smith was supported by a National Science Foundation Graduate Research Fellowship. This work was supported by the National Institutes of Health (GM100233 and HG012287), the John Templeton Foundation (grant 61220), and by the Allen Discovery Center program, a Paul G. Allen Frontiers Group advised program of the Paul G. Allen Family Foundation and the Howard Hughes Medical Institute.

## Author Information

### Contributions

A.H., D.R. and V.M.N. supervised the study. M.K.L, O.S. and A.A. analyzed genetic data. M.K.L., A.H., D.R. and V.M.N. wrote the manuscript with input from all co-authors.

## Supplementary Information

**Supplementary Table 1**

This file contains metadata information for all samples used in the analysis.

**Supplementary Table 2**

This file contains the significant gene sets from enrichment analysis for the Neolithic period.

**Supplementary Table 3**

This file contains the significant gene sets from enrichment analysis for the Bronze Age period.

**Supplementary Table 4**

This file contains the significant gene sets from enrichment analysis for the Historical period.

**Supplementary Table 5**

This file contains the results of regressing the effect sizes of traits from the Biobank of Japan on PC loadings.

**Supplementary Table 6**

This file contains the results of regressing the effect sizes of traits from the UK Biobank on PC loadings.

**Supplementary Table 7**

This file contains the results of the analysis on the correlations and shared effect directions for effect sizes of matched traits in the Biobank of Japan and the UK Biobank.

**Supplementary Table 8**

This file contains the results for the polygenic selection scan using the directions of effect sizes for traits from the Biobank of Japan.

**Supplementary Table 9**

This file contains the resampling bin distributions of null variants for the polygenic selection scan on a single trait.

**Supplementary Table 10**

This file contains the results for the polygenic selection scan using both the magnitudes and directions of effect sizes for traits from the Biobank of Japan.

**Supplementary Table 11**

This file contains the results for the polygenic selection scan using the directions of effect sizes for traits from the Biobank of Japan and removing SNPs that were in the lowest 27.5% of the chi-squared statistic distribution.

**Supplementary Table 12**

This file contains the results for the polygenic selection scan using the directions of effect sizes for traits from the UK Biobank.

**Supplementary Table 13**

This file contains the results for the polygenic selection scan using the directions of effect sizes for traits from the within-sibling GWAS consortium.

**Supplementary Data 1**

This file contains genome-wide selection scan results and allele frequencies for the Neolithic period.

**Supplementary Data 2**

This file contains genome-wide selection scan results and allele frequencies for the Bronze Age period.

**Supplementary Data 3**

This file contains genome-wide selection scan results and allele frequencies for the Historical period.

## References

1. Richerson, P. J., Boyd, R. & Henrich, J. Gene-culture coevolution in the age of genomics. Proc Natl Acad Sci U S A 107, 8985–8992 (2010).

2. Richerson, P. J. & Boyd, R. Not by Genes Alone : How Culture Transformed Human Evolution. (University of Chicago Press, 2008).

3. Armelagos, G. & Cohen, M. Paleopathology at the Origins of Agriculture. (1984).

4. Mathieson, I. et al. Genome-wide patterns of selection in 230 ancient Eurasians. Nature 2015 528:7583 528, 499–503 (2015).

5. Margaryan, A. et al. Population genomics of the Viking world. Nature 2020 585:7825 585, 390–396 (2020).

6. Marnetto, D. et al. Ancestral genomic contributions to complex traits in contemporary Europeans. Current Biology 32, 1412-1419.e3 (2022).

7. Mathieson, S. & Mathieson, I. FADS1 and the Timing of Human Adaptation to Agriculture. Molecular Biology and Evolution 35, 2957–2970 (2018).

8. Segurel, L. et al. Why and when was lactase persistence selected for? Insights from Central Asian herders and ancient DNA. PLOS Biology 18, e3000742 (2020).

9. Ju, D. & Mathieson, I. The evolution of skin pigmentation-associated variation in West Eurasia. Proc Natl Acad Sci U S A 118, (2020).

10. Domínguez-Andrés, J. et al. Evolution of cytokine production capacity in ancient and modern european populations. Elife 10, (2021).

11. Allentoft, M. E. et al. Population Genomics of Stone Age Eurasia. bioRxiv 2022.05.04.490594 (2022) doi:10.1101/2022.05.04.490594.

12. Kerner, G. et al. Genetic adaptation to pathogens and increased risk of inflammatory disorders in post-Neolithic Europe. bioRxiv 2022.07.02.498543 (2022) doi:10.1101/2022.07.02.498543.

13. Mathieson, I. & Terhorst, J. Direct detection of natural selection in Bronze Age Britain. bioRxiv 2022.03.14.484330 (2022) doi:10.1101/2022.03.14.484330.

14. Souilmi, Y. et al. Admixture has obscured signals of historical hard sweeps in humans. bioRxiv 2020.04.01.021006 (2021) doi:10.1101/2020.04.01.021006.

15. Childebayeva, A. et al. Population Genetics and Signatures of Selection in Early Neolithic European Farmers. Molecular Biology and Evolution 39, (2022).

16. Mühlemann, B. et al. Diverse variola virus (smallpox) strains were widespread in northern Europe in the Viking Age. Science 369, (2020).

17. Spyrou, M. A. et al. Phylogeography of the second plague pandemic revealed through analysis of historical Yersinia pestis genomes. Nature Communications 2019 10:1 10, 1–13 (2019).

18. Bos, K. I. et al. Eighteenth century Yersinia pestis genomes reveal the long-term persistence of an historical plague focus. Elife 5, (2016).

19. Fu, Q. et al. An early modern human from Romania with a recent Neanderthal ancestor. Nature 524, 216 (2015).

20. Rohland, N. et al. Three Reagents for in-Solution Enrichment of Ancient Human DNA at More than a Million SNPs. bioRxiv 2022.01.13.476259 (2022) doi:10.1101/2022.01.13.476259.

21. Rubinacci, S., Ribeiro, D. M., Hofmeister, R. J. & Delaneau, O. Efficient phasing and imputation of low-coverage sequencing data using large reference panels. Nature Genetics 2021 53:1 53, 120–126 (2021).

22. Haak, W. et al. Massive migration from the steppe was a source for Indo-European languages in Europe. Nature 2015 522:7555 522, 207–211 (2015).

23. Lazaridis, I. et al. Genomic insights into the origin of farming in the ancient Near East. Nature 2016 536:7617 536, 419–424 (2016).

24. Allentoft, M. E. et al. Population genomics of Bronze Age Eurasia. Nature 2015 522:7555 522, 167–172 (2015).

25. Papac, L. et al. Dynamic changes in genomic and social structures in third millennium BCE central Europe. Science Advances 7, 6941–6966 (2021).

26. Lipson, M. et al. Parallel palaeogenomic transects reveal complex genetic history of early European farmers. Nature 551, 368–372 (2017).

27. Narasimhan, V. M. et al. The Formation of Human Populations in South and Central Asia. Science 365, (2019).

28. Olalde, I. et al. The Beaker phenomenon and the genomic transformation of northwest Europe. Nature 2018 555:7695 555, 190–196 (2018).

29. Olalde, I. et al. The genomic history of the Iberian Peninsula over the past 8000 years. Science (1979) 363, 1230–1234 (2019).

30. Risch, N. & Merikangas, K. The future of genetic studies of complex human diseases. Science 273, 1516–1517 (1996).

31. Devlin, B. & Roeder, K. Genomic control for association studies. Biometrics 55, 997–1004 (1999).

32. Cuadros-Espinoza, S., Laval, G., Quintana-Murci, L. & Patin, E. The genomic signatures of natural selection in admixed human populations. The American Journal of Human Genetics 109, 710–726 (2022).

33. Fauman, E. B. & Hyde, C. An optimal variant to gene distance window derived from an empirical definition of cis and trans protein QTLs. BMC Bioinformatics 23, 169 (2022).

34. Watanabe, K., Taskesen, E., van Bochoven, A. & Posthuma, D. Functional mapping and annotation of genetic associations with FUMA. Nature Communications 2017 8:1 8, 1–11 (2017).

35. Claussnitzer, M. et al. FTO Obesity Variant Circuitry and Adipocyte Browning in Humans. New England Journal of Medicine 373, 895–907 (2015).

36. Smemo, S. et al. Obesity-associated variants within FTO form long-range functional connections with IRX3. Nature 507, 371 (2014).

37. Aguet, F. et al. The GTEx Consortium atlas of genetic regulatory effects across human tissues. Science (1979) 369, 1318–1330 (2020).

38. Lee, N. K. et al. Endocrine Regulation of Energy Metabolism by the Skeleton. Cell 130, 456–469 (2007).

39. Martin Carli, J. F., LeDuc, C. A., Zhang, Y., Stratigopoulos, G. & Leibel, R. L. FTO mediates cell-autonomous effects on adipogenesis and adipocyte lipid content by regulating gene expression via 6mA DNA modifications. Journal of Lipid Research 59, 1446–1460 (2018).

40. Matsuoka, T. et al. Members of the large Maf transcription family regulate insulin gene transcription in islet beta cells. Mol Cell Biol 23, 6049–6062 (2003).

41. Daily, J. W. & Park, S. Interaction of BDNF rs6265 variants and energy and protein intake in the risk for glucose intolerance and type 2 diabetes in middle-aged adults. Nutrition 33, 187–194 (2017).

42. Bathina, S. & Das, U. N. Brain-derived neurotrophic factor and its clinical implications. Archives of Medical Science : AMS 11, 1164 (2015).

43. Lindesmith, L. C. et al. Mechanisms of GII.4 Norovirus Persistence in Human Populations. PLoS Medicine 5, 0269–0290 (2008).

44. Hazra, A. et al. Common variants of FUT2 are associated with plasma vitamin B12 levels. Nat Genet 40, 1160 (2008).

45. Peters, V. A., Joesting, J. J. & Freund, G. G. IL-1 receptor 2 (IL-1R2) and its role in immune regulation. Brain Behav Immun 32, 1 (2013).

46. Thali, M. et al. Functional association of cyclophilin A with HIV-1 virions. Nature 372, 363–365 (1994).

47. Erdogmus, S. et al. Cavβ1 regulates T cell expansion and apoptosis independently of voltage-gated Ca2+ channel function. Nature Communications 2022 13:1 13, 1–19 (2022).

48. Yuki, K. E. et al. CYRI/FAM49B negatively regulates RAC1-driven cytoskeletal remodelling and protects against bacterial infection. Nat Microbiol 4, 1516–1531 (2019).

49. Shang, W. et al. Genome-wide CRISPR screen identifies FAM49B as a key regulator of actin dynamics and T cell activation. Proc Natl Acad Sci U S A 115, E4051–E4060 (2018).

50. Key, F. M. et al. Emergence of human-adapted Salmonella enterica is linked to the Neolithization process. Nature Ecology & Evolution 2020 4:3 4, 324–333 (2020).

51. Ju, D. & Mathieson, I. The evolution of skin pigmentation-associated variation in West Eurasia. Proc Natl Acad Sci U S A 118, (2020).

52. Field, Y. et al. Detection of human adaptation during the past 2000 years. Science 354, 760 (2016).

53. Harriff, M. J. et al. MR1 displays the microbial metabolome driving selective MR1-restricted T cell receptor usage. Science Immunology 3, 2556 (2018).

54. Augusto, D. G., Norman, P. J., Dandekar, R. & Hollenbach, J. A. Fluctuating and geographically specific selection characterize rapid evolution of the human Kir region. Frontiers in Immunology 10, 989 (2019).

55. Cao, C. et al. Structure, function and pharmacology of human itch GPCRs. Nature 2021 600:7887 600, 170–175 (2021).

56. McNeil, B. D. et al. Identification of a mast-cell-specific receptor crucial for pseudo-allergic drug reactions. Nature 519, 237–241 (2015).

57. Wierbowski, S. D. et al. A 3D structural SARS-CoV-2–human interactome to explore genetic and drug perturbations. Nature Methods 2021 18:12 18, 1477–1488 (2021).

58. Pietzner, M. et al. Genetic architecture of host proteins involved in SARS-CoV-2 infection. Nature Communications 2020 11:1 11, 1–14 (2020).

59. Merad, M. & Martin, J. C. Author Correction: Pathological inflammation in patients with COVID-19: a key role for monocytes and macrophages (Nature Reviews Immunology, (2020), 20, 6, (355-362), 10.1038/s41577-020-0331-4). Nature Reviews Immunology 20, 448 (2020).

60. Rascovan, N. et al. Emergence and Spread of Basal Lineages of Yersinia pestis during the Neolithic Decline. Cell 176, 295-305.e10 (2019).

61. Andrades Valtueña, A. et al. Stone Age Yersinia pestis genomes shed light on the early evolution, diversity, and ecology of plague. Proceedings of the National Academy of Sciences 119, (2022).

62. Lu, H., Cassis, L. A., Kooi, C. W. V. & Daugherty, A. Structure and functions of angiotensinogen. Hypertension Research 2016 39:7 39, 492–500 (2016).

63. Kennedy, M. A. et al. ABCG1 has a critical role in mediating cholesterol efflux to HDL and preventing cellular lipid accumulation. Cell Metab 1, 121–131 (2005).

64. Olbrich, H. et al. Recessive HYDIN Mutations Cause Primary Ciliary Dyskinesia without Randomization of Left-Right Body Asymmetry. The American Journal of Human Genetics 91, 672–684 (2012).

65. Patterson, N. et al. Large-scale migration into Britain during the Middle to Late Bronze Age. Nature 2021 601:7894 601, 588–594 (2021).

66. Jablonski, N. G. & Chaplin, G. Human skin pigmentation as an adaptation to UV radiation. Proc Natl Acad Sci U S A 107, 8962–8968 (2010).

67. Albiñana, C. et al. Genetic correlates of vitamin D-binding protein and 25 hydroxyvitamin D in neonatal dried blood spots. medRxiv 2022.06.08.22276164 (2022) doi:10.1101/2022.06.08.22276164.

68. Zhernakova, A. et al. Evolutionary and functional analysis of celiac risk loci reveals SH2B3 as a protective factor against bacterial infection. Am J Hum Genet 86, 970–977 (2010).

69. Hernandez, R. D. et al. Classic selective sweeps were rare in recent human evolution. Science (1979) 331, 920–924 (2011).

70. Murphy, D., Elyashiv, E., Amster, G. & Sella, G. Broad-scale variation in human genetic diversity levels is predicted by purifying selection on coding and non-coding elements. bioRxiv 2021.07.02.450762 (2021) doi:10.1101/2021.07.02.450762.

71. Sella, G. & Barton, N. H. Thinking About the Evolution of Complex Traits in the Era of Genome-Wide Association Studies. https://doi.org/10.1146/annurev-genom-083115-022316 20, p461–493 (2019).

72. Berg, J. J. & Coop, G. A Population Genetic Signal of Polygenic Adaptation. PLOS Genetics 10, e1004412 (2014).

73. Höllinger, I., Pennings, P. S. & Hermisson, J. Polygenic adaptation: From sweeps to subtle frequency shifts. PLOS Genetics 15, e1008035 (2019).

74. Speidel, L., Forest, M., Shi, S. & Myers, S. R. A method for genome-wide genealogy estimation for thousands of samples. Nature Genetics 2019 51:9 51, 1321–1329 (2019).

75. Berg, J. J. et al. Reduced signal for polygenic adaptation of height in UK biobank. Elife 8, (2019).

76. Sohail, M. et al. Polygenic adaptation on height is overestimated due to uncorrected stratification in genome-wide association studies. Elife 8, (2019).

77. Rosenberg, N. A., Edge, M. D., Pritchard, J. K. & Feldman, M. W. Interpreting polygenic scores, polygenic adaptation, and human phenotypic differences. Evolution, Medicine, and Public Health 2019, 26–34 (2019).

78. Harpak, A. & Przeworski, M. The evolution of group differences in changing environments. PLOS Biology 19, e3001072 (2021).

79. Mills, M. C. & Mathieson, I. The challenge of detecting recent natural selection in human populations. Proceedings of the National Academy of Sciences 119, (2022).

80. Young, A. I., Benonisdottir, S., Przeworski, M. & Kong, A. Deconstructing the sources of genotype-phenotype associations in humans. Science (1979) 365, 1396–1400 (2019).

81. Mostafavi, H. et al. Variable prediction accuracy of polygenic scores within an ancestry group. Elife 9, (2020).

82. Barton, N., Hermisson, J. & Nordborg, M. Population Genetics: Why structure matters. Elife 8, (2019).

83. Chen, M. et al. Evidence of Polygenic Adaptation in Sardinia at Height-Associated Loci Ascertained from the Biobank Japan. American Journal of Human Genetics 107, 60–71 (2020).

84. Zhou, W. et al. Global Biobank Meta-analysis Initiative: powering genetic discovery across human diseases. Unnur Thorsteinsdottir 27,.

85. Kanai, M. et al. Insights from complex trait fine-mapping across diverse populations. medRxiv 2021.09.03.21262975 (2021) doi:10.1101/2021.09.03.21262975.

86. des Marais, D. L., Hernandez, K. M. & Juenger, T. E. Genotype-by-environment interaction and plasticity: Exploring genomic responses of plants to the abiotic environment. Annual Review of Ecology, Evolution, and Systematics 44, 5–29 (2013).

87. Halldorsson, B. v. et al. Characterizing mutagenic effects of recombination through a sequence-level genetic map. Science 363, (2019).

88. McVicker, G., Gordon, D., Davis, C. & Green, P. Widespread Genomic Signatures of Natural Selection in Hominid Evolution. PLOS Genetics 5, e1000471 (2009).

89. Cox, S. L., Ruff, C. B., Maier, R. M. & Mathieson, I. Genetic contributions to variation in human stature in prehistoric Europe. Proc Natl Acad Sci U S A 116, 21484–21492 (2019).

90. Howe, L. J. et al. Within-sibship genome-wide association analyses decrease bias in estimates of direct genetic effects. Nature Genetics 2022 54:5 54, 581–592 (2022).

91. Pybus, M. et al. Hierarchical boosting: a machine-learning framework to detect and classify hard selective sweeps in human populations. Bioinformatics 31, 3946–3952 (2015).

92. Maier, R. et al. No statistical evidence for an effect of CCR5-Δ32 on lifespan in the UK Biobank cohort. Nature Medicine 2019 26:2 26, 178–180 (2019).

93. Wei, X. & Nielsen, R. Retraction Note: CCR5-Δ32 is deleterious in the homozygous state in humans. Nature Medicine 2019 25:11 25, 1796–1796 (2019).

94. Fortier, A. L. & Pritchard, J. K. Ancient Trans-Species Polymorphism at the Major Histocompatibility Complex in Primates. bioRxiv 2022.06.28.497781 (2022) doi:10.1101/2022.06.28.497781.

95. Marciniak, S. et al. An integrative skeletal and paleogenomic analysis of prehistoric stature variation suggests relatively reduced health for early European farmers. bioRxiv 12, 2021.03.31.437881 (2021).

96. Shennan, S. et al. Regional population collapse followed initial agriculture booms in mid-Holocene Europe. Nature Communications 2013 4:1 4, 1–8 (2013).

97. Evershed, R. P. et al. Dairying, diseases and the evolution of lactase persistence in Europe. Nature 2022 1–10 (2022) doi:10.1038/s41586-022-05010-7.

98. de Barros Damgaard, P. et al. 137 ancient human genomes from across the Eurasian steppes. Nature 557, 369–374 (2018).

99. Mühlemann, B. et al. Ancient hepatitis B viruses from the Bronze Age to the Medieval period. Nature 2018 557:7705 557, 418–423 (2018).

100. Guellil, M. et al. Ancient herpes simplex 1 genomes reveal recent viral structure in Eurasia. Science Advances 8, 4435 (2022).

101. Key, F. M. et al. Emergence of human-adapted Salmonella enterica is linked to the Neolithization process. Nature Ecology & Evolution 2020 4:3 4, 324–333 (2020).

102. Kocher, A. et al. Ten millennia of hepatitis B virus evolution. Science (1979) 374, (2021).

103. Verdugo, M. P. et al. Ancient cattle genomics, origins, and rapid turnover in the Fertile Crescent. Science (1979) 365, 173–176 (2019).

104. Librado, P. et al. The origins and spread of domestic horses from the Western Eurasian steppes. Nature 2021 598:7882 598, 634–640 (2021).

105. Chen, L. et al. Inflammatory responses and inflammation-associated diseases in organs. Oncotarget 9, 7204 (2018).

106. Church, D. M. et al. Modernizing reference genome assemblies. PLoS Biol 9, (2011).

107. Li, H. & Durbin, R. Fast and accurate short read alignment with Burrows-Wheeler transform. Bioinformatics 25, 1754–1760 (2009).

108. Fu, Q. et al. A Revised Timescale for Human Evolution Based on Ancient Mitochondrial Genomes. Current Biology 23, 553–559 (2013).

109. Korneliussen, T. S., Albrechtsen, A. & Nielsen, R. ANGSD: Analysis of Next Generation Sequencing Data. BMC Bioinformatics 15, 1–13 (2014).

110. Narasimhan, V. et al. BCFtools/RoH: a hidden Markov model approach for detecting autozygosity from next-generation sequencing data. Bioinformatics 32, 1749 (2016).

111. Patterson, N., Price, A. L. & Reich, D. Population Structure and Eigenanalysis. PLOS Genetics 2, e190 (2006).

112. Lazaridis, I. et al. Ancient human genomes suggest three ancestral populations for present-day Europeans. Nature 2014 513:7518 513, 409–413 (2014).

113. Patterson, N. et al. Ancient admixture in human history. Genetics 192, 1065–1093 (2012).

114. Reich, D., Thangaraj, K., Patterson, N., Price, A. L. & Singh, L. Reconstructing Indian population history. Nature 2009 461:7263 461, 489–494 (2009).

115. scipy.optimize.minimize — SciPy v1.8.1 Manual. https://docs.scipy.org/doc/scipy/reference/generated/scipy.optimize.minimize.html#scipy.optimize.minimize.

116. McLaren, W. et al. The Ensembl Variant Effect Predictor. Genome Biology 17, 1–14 (2016).

117. Boughton, A. P. et al. LocusZoom.js: Interactive and embeddable visualization of genetic association study results. Bioinformatics 37, 3017–3018 (2021).

118. Choin, J. et al. Genomic insights into population history and biological adaptation in Oceania. Nature 2021 592:7855 592, 583–589 (2021).

119. UK Biobank — Neale lab. http://www.nealelab.is/uk-biobank/.

120. Sakaue, S. et al. A global atlas of genetic associations of 220 deep phenotypes. medRxiv 46, 2020.10.23.20213652 (2021).

121. McVicker, G., Gordon, D., Davis, C. & Green, P. Widespread Genomic Signatures of Natural Selection in Hominid Evolution. PLOS Genetics 5, e1000471 (2009).

